# *Xanthomonas oryzae* pv. *oryzae* type-III effector TAL9b targets a broadly conserved disease susceptibility locus to promote pathogenesis in rice

**DOI:** 10.1101/2024.05.01.592040

**Authors:** C.G. Gokulan, Sohini Deb, Namami Gaur, Apoorva Masade, Niranjan Gattu, P.R. Rennya, Nisha Sao, Donald James, Ramesh V. Sonti, Hitendra K. Patel

**Author notes:** Department of Plant and Environmental Sciences, Section for Plant and Soil Sciences, University of Copenhagen, Thorvaldsensvej 40, 1871 Frederiksberg C, Denmark (Sohini Deb). The James Hutton Institute, Invergowrie, DD2 5DA, Scotland, UK. Kerala Forest Research Institute, Peechi, India - 680653 (Donald James). Equally contributed authors (C.G. Gokulan, Sohini Deb, Namami Gaur). Corresponding authors **Addresses for correspondence:** Ramesh V. Sonti; Hitendra K Patel.

## Abstract

*Xanthomonas oryzae* pv. *oryzae* (Xoo), the causal agent of bacterial blight of rice, translocates multiple Transcription Activator-Like Effectors (TALEs) into rice cells. The TALEs localize to the host cell nucleus, where they bind to the DNA in a sequence-specific manner and enhance gene expression to promote disease susceptibility. Xoo strain PXO99^A^ encodes nineteen TALEs, but the host targets of all these TALEs have not been defined. A meta-analysis of rice transcriptome profiles revealed a gene annotated as flavonol synthase/flavanone-3 hydroxylase (henceforth *OsS5H/FNS-03g*) to be highly induced upon Xoo infection. Further analyses revealed that this gene is induced by PXO99^A^ using TAL9b, a broadly conserved TALE of Xoo. Disruption of *tal9b* rendered PXO99^A^ less virulent. OsS5H/FNS-03g functionally complemented its *Arabidopsis* homologue AtDMR6, a well-studied disease susceptibility locus. Biochemical analyses suggested that OsS5H/FNS-03g is a bifunctional protein with Salicylic Acid 5’ Hydroxylase (S5H) and Flavone Synthase-I (FNS-I) activities. Further, an exogenous application of apigenin on rice leaves, the flavone that is enzymatically produced by OsS5H/FNS-03g, promoted virulence of PXO99^A^ *tal9b*-. Overall, our study suggests that OsS5H/FNS-03g is a bifunctional enzyme and its product apigenin is potentially involved in promoting Xoo virulence.

## INTRODUCTION

Phytopathogens deploy several virulence factors to control host physiology and promote pathogenesis (Wang and Wang, 2018). The genus *Xanthomonas* comprises several economically important phytopathogens that cause severe yield losses in multiple important food crops globally (White et al., 2009). *Oryza sativa* (henceforth rice) is an important food crop that feeds about half of the global human population (Fukagawa and Ziska, 2019). Rice production is hampered by multiple diseases, one of most devastating is the bacterial blight (BB) disease caused by *Xanthomonas oryzae* pv. *oryzae* (Xoo; Savary et al., 2019). Yet another bacterial disease of rice is bacterial leaf streak (BLS) caused by *X. oryzae* pv. *oryzicola* (Xoc). Although Xoo and Xoc are pathovars of *X. oryzae,* their mode of infection and colonization in rice leaves are different. Xoo, a vascular pathogen, enters through either hydathodes or wounds and colonizes the xylem vessels. On the other hand, Xoc invades rice through stomata or wounds and colonizes the mesophyll tissue (Niño-Liu et al., 2006). Both Xoo and Xoc utilize the type III secretion system (T3SS) to deploy a specific class of effectors known as Transcription Activator-Like Effectors (TALEs) directly into the rice cells to promote pathogenesis (White et al., 2009).

TALEs consist of ∼34 conserved repeating amino acids that vary at positions 12 and 13 in each repeat. These pairs of residues called Repeat Variable Diresidue (RVD) interact with the Effector Binding Element (EBE) in the promoters of host target genes and modulate their expression (Boch et al., 2009; Bogdanove et al., 2010). More often than not, the TAL effector targeted host genes confer disease susceptibility and are known as susceptibility factors (Antony et al., 2010; Cernadas et al., 2014; Peng et al., 2019; Streubel et al., 2013; Sugio et al., 2007; Tran et al., 2018; Wu et al., 2022; Yang et al., 2006). Understanding the mechanisms of TALE-mediated host susceptibility is important to devise strategies for disease resistance breeding. For instance, certain Xoo-delivered TALEs recognize EBE in the genes encoding the SUGARS WILL EVENTUALLY BE EXPORTED TRANSPORTER (SWEET) family proteins and induce their expression to enhance apoplastic sucrose availability (Bezrutczyk et al., 2018; Streubel et al., 2013). Interfering with the ability of Xoo to induce *SWEET* genes makes the plants resistant to Xoo infection (Blanvillain-Baufumé et al., 2017; Chu et al., 2006; Oliva et al., 2019). Further, Mücke et al., (2019) identified several rice genes that are directly induced by Xoo TALEs. Besides the examples in Xoo, studies have also been conducted on the role of TALEs in Xoc pathogenesis. For instance, severe water-soaked leaf streaking was shown to be associated with Tal2g_BLS256_-dependent upregulation of a rice gene called *OsSULTR3;6* (Cernadas et al., 2014). Another study reported that Tal7_RS105_ activates the promoters of two rice genes *Os09g29100* (predicted to encode Cyclin-D4-1) and *Os12g42970* (predicted to encode a GATA zinc finger family protein), and that Tal7_RS105_ suppresses avrXa7-Xa7 mediated defense in Rice (Cai et al., 2017).

A recent study identified a rice FLAVANONE 3’ HYDROXYLASE encoding gene (*OsF3H_03g’_ LOC_Os03g03034*) to be targeted by Xoc in a TALE-dependent manner. Notably, the presence of the cognate TALE that induces *OsF3H_03g_* correlated with the hypervirulence of the tested Xoc strain (Wu et al., 2022). Earlier it was reported that the same gene (called *OsFNS* therein) is induced by Xoo TALEs that belong to the class TAL AQ (that includes TAL9b in Xoo PXO99^A^) by introducing the TALE into an Xoo strain that lacks TAL effectors (Mücke et al., 2019). Intriguingly, *OsF3H_03g_* is one of the very few genes that is induced by TALEs from Xoo as well as Xoc (Cernadas et al., 2014; Mücke et al., 2019). *OsF3H_03g_* has been annotated with various names in annotation servers and by different studies including 2-oxoglutarate dependent dioxygenase (2-ODD/DOX), flavone synthase (FNS), flavanone 3’ hydroxylase (F3H), and salicylic acid (SA) 5’ hydroxylase (S5H). Two recent studies reported that OsF3H_03g_ catalyses SA 5’ hydroxylation to form 2,5-dihydroxybenzoic acid (2,5-DHBA) and established that mutating rice S5Hs resulted in broad-spectrum disease resistance (Liu et al., 2023; Zhang et al., 2022). On the other hand, an earlier study showed the FNS activity of OsF3H_03g_ (Kim et al., 2008). However, no study has formally shown that purified OsF3H_03g_ is a bifunctional protein that possesses both FNS and S5H biochemical activities. This lack of clarity on the activity of OsF3H_03g_ has been previously highlighted (Mücke et al., 2019; Wu et al., 2022).

Here, we validated the predicted role of TAL9b in inducing *OsF3H_03g_* and showed that the disruption of *tal9b* compromised the virulence of PXO99^A^. Ectopic expression of *OsF3H_03g_* in *Arabidopsis atdmr6* mutant plants restored susceptibility of the plants to *Pseudomonas syringae* pv. *tomato* DC3000, indicating that OsF3H_03g_ is a functional homologue of *Arabidopsis* DOWNY MILDEW RESISTANT 6 (DMR6). Biochemical assays using purified recombinant protein indicated that the protein performs dual functions *in vitro* i.e., SA hydroxylation and flavone synthesis. Therefore, we address the protein henceforth as OsS5H/FNS-03g. Furthermore, we identified a virulence promoting activity of the flavone apigenin - enzymatically produced by OsS5H/FNS-03g - wherein it enhanced the virulence of the PXO99^A^ tal9b-strain. Overall, our study expands the bacterial blight susceptibility gene repertoire of rice and reinforces that DMR6 homologues are an evolutionarily conserved plant disease susceptibility hub and are a potential target for disease resistance breeding in various crops.

## RESULTS

### The rice gene *LOC_Os03g03034* is induced by multiple pathogens and pests

Analysis of publicly available gene expression data (Table S1) revealed that the rice gene *LOC_Os03g03034* is highly induced by *Xanthomonas oryzae* pv. *oryzae, Xanthomonas oryzae* pv. *oryzicola, Magnaporthe oryzae,* and infestation by the brown plant hopper (*Nilaparvata lugens*) insect pest but not by a non-host pathogen *Puccinia triticina* f. sp. *tritici* (Figure 1a). Semi-quantitative PCR analyses indicated that the transcript variant *LOC_Os03g03034.1* is primarily induced by the tested strains (Figure S1) and thus has been used throughout this study. Data mining from a public RNA-seq database revealed that the expression of *OsS5H/FNS-03g* is induced primarily in pathogenic interactions as compared to beneficial interactions (Figure S2a). Notably, induction of the gene, on average, was higher during shoot/foliar pathogenesis (Figure S2b). We found that multiple strains of the *X. oryzae* pathovars considerably induce the expression of *OsS5H/FNS-03g* (Figure 1b). We further observed that the expression of *OsS5H/FNS-03g* increases temporally during Xoo infection of rice (Figure S3a). Previous studies have reported that *OsS5H/FNS-03g* could be a target of Xoo and Xoc type III-secreted TAL effectors. To validate the prediction, we generated the mutants of Xoo PXO99^A^ and Xoc BXOR1 that are defective in the type III secretion system (T3SS). Treatment of rice leaves with wild-type and T3SS mutant strains of both Xoo and Xoc showed that the induction of *OsS5H/FNS-03g* is dependent on functional T3SS (Figure 1c). Taken together, the results suggest that *OsS5H/FNS-03g* is a potential susceptibility gene to multiple shoot/foliar pathogens and pests of rice.

**Figure 1:**
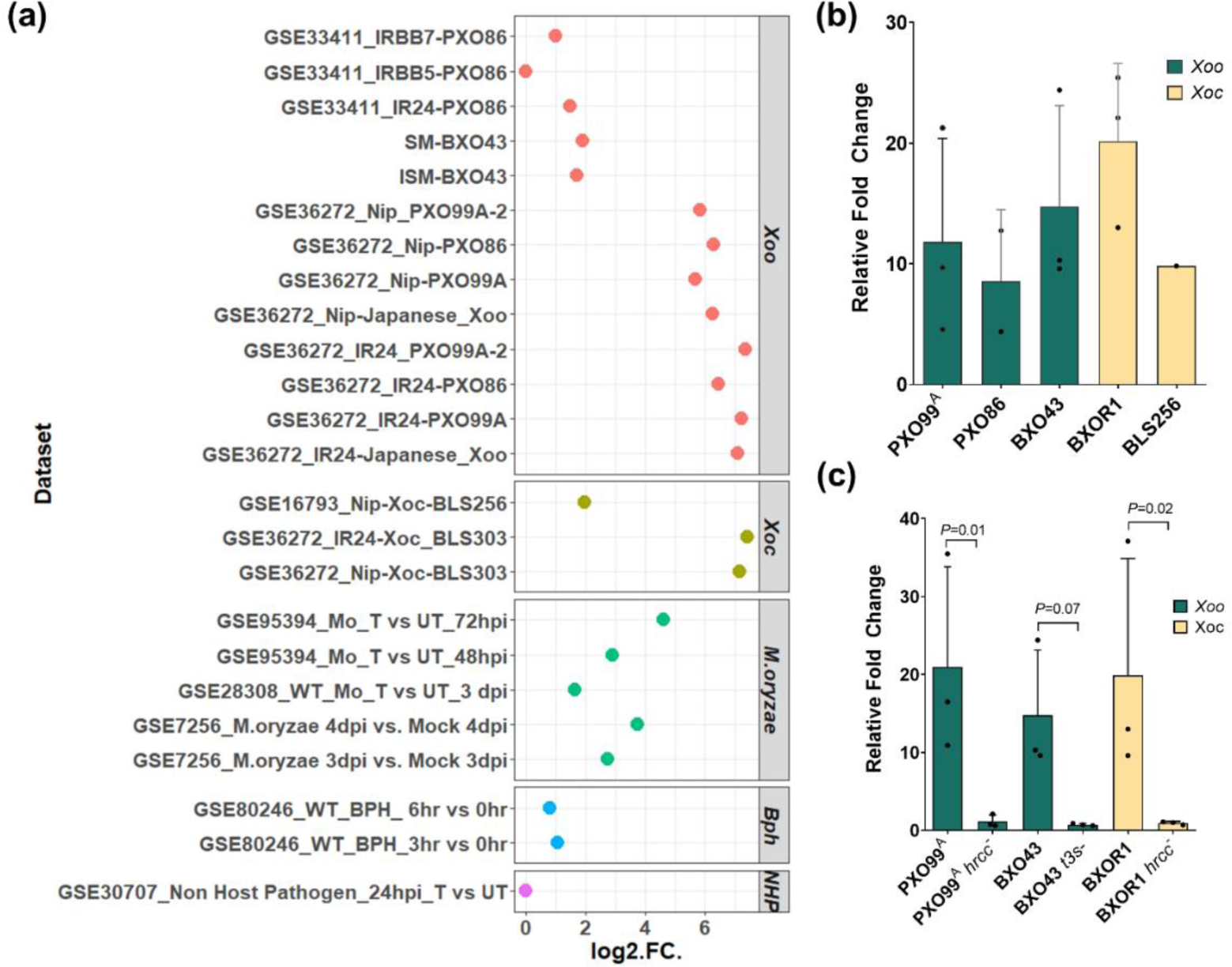
Expression of *OsS5H/FNS-03g* is highly induced during infection by various pathogens and a pest. (a) Various publicly available transcriptomics data were analysed and *OsS5H/FNS-03g* was observed to be highly induced in pathogenic interactions of rice. Publicly available microarray data were retrieved from NCBI-Gene Expression Omnibus and analysed further to obtain the fold change values upon treatment as compared to the corresponding mock treated samples. (b, c) Bacterial suspensions of indicated strains were infiltrated into the leaves of 14-day old rice leaves, sampled at 24 hours post infiltration and the gene expression was analysed using qRT-PCR. The gene expression levels were normalised with the internal control *OsGAPDH*. Fold change values were calculated using the 2^-ΔΔCt^ method relative to the mock-infiltrated samples. Error bars in (b) and (c) indicate the standard deviation of the fold change values in a minimum of two independent experiments. *P* values were calculated using the One-way ANOVA test.

### Disruption of TAL9b affects *OsS5H/FNS-03g* induction and virulence of Xoo PXO99^A^

Previous reports indicated the presence of an EBE in the promoter of *OsS5H/FNS-03g* (Cernadas et al., 2014; Mücke et al., 2019); Figure S4). Intriguingly, effectors from both the pathovars of *X. oryzae* i.e., Xoo, and Xoc, utilise TALEs of different families to induce the expression of *OsS5H/FNS-03g* while binding to the same EBE (Mücke et al., 2019; Figure S5). *In silico* prediction analysis indicated that TAL9b encoded by Xoo PXO99^A^ is likely the cognate TALE that activates the expression of *OsS5H/FNS-03g.* Therefore, a Tal9b-defective mutant of Xoo PXO99^A^ (PXO99^A^ tal9b-) was generated through insertional mutagenesis to study the role of TAL9b in *OsS5H/FNS-03g* induction and virulence of Xoo. Treatment of rice leaves with PXO99^A^ *tal9b*-exhibited a reduced induction of *OsS5H/FNS-03g* transcripts when compared to induction by the wild-type strain (Figure 2a). Further, the Tal9b mutant showed significantly reduced virulence when compared to the wild-type strain on a susceptible rice line (Figure 2b, c). These results indicate that optimal induction of *OsS5H/FNS-03g* by TAL9b is required for full virulence of Xoo PXO99^A^.

**Figure 2:**
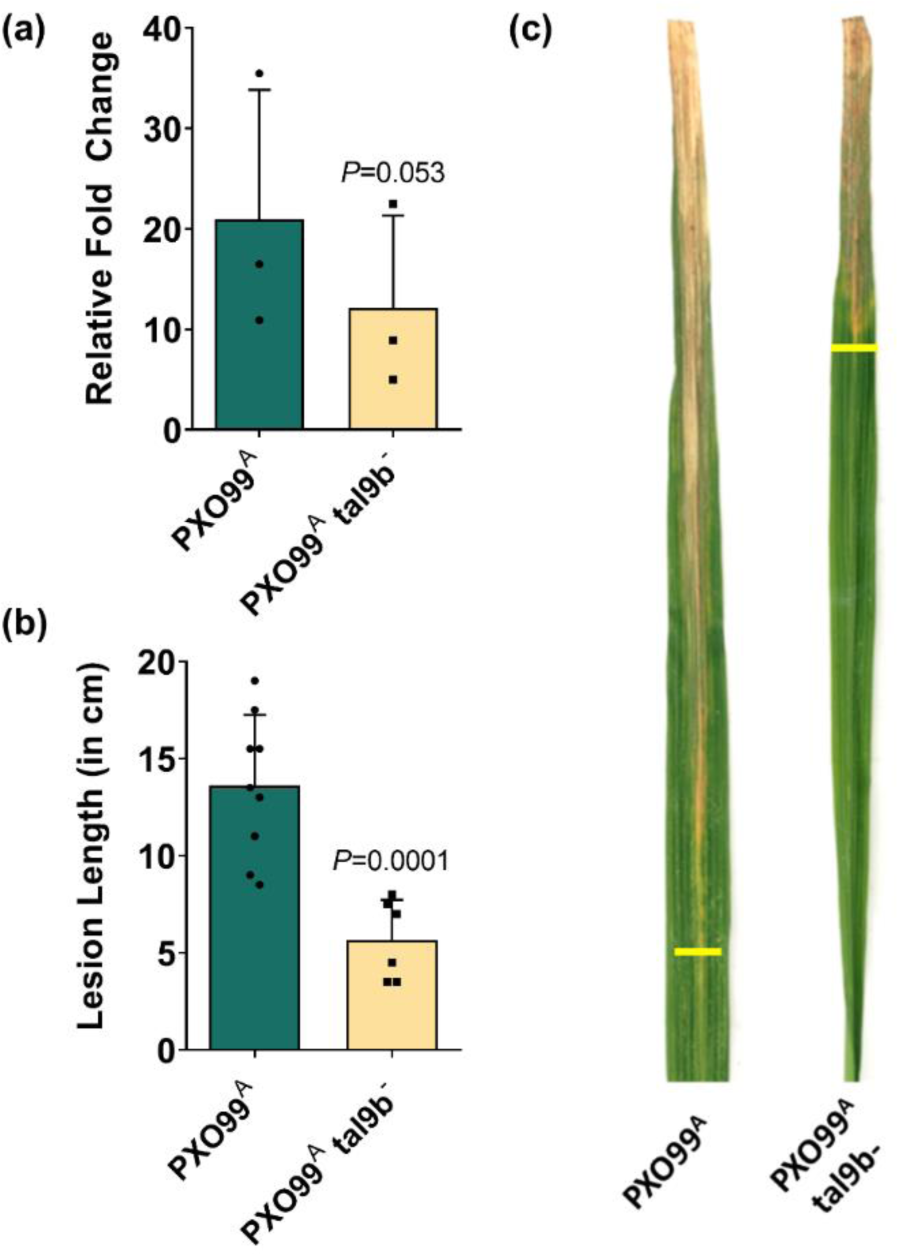
Tal9b is required for induction of *OsS5H/FNS-03G* and full virulence of Xoo PXO99^A^. (a) Gene expression analysis using qRT-PCR showed a reduced induction *of OsS5H/FNS-03g* transcripts by the *Tal9b* mutant of Xoo PXO99^A^. The gene expression quantification was performed from leaf samples at 24 hours post infiltrating a suspension of the described Xoo strains of OD_600nm_=1 into the leaves of 14-day old TN1 rice plants. Error bars indicate the standard deviation of the fold change values obtained with respect to the mock-treated samples in three independent experiments. *P*-value was calculated using a two-tailed paired *t* test. (b) Rice plants when 60 days old, were clip inoculated with PXO99^A^ and PXO99^A^ *tal9b*-strains of OD_600nm_=1. Bars in (b) indicate the average lesion length (n≥6) at 14-days post infection (dpi) and the error bars represent the standard deviation. The *P*-value was calculated using a two-tailed unpaired *t* test. Similar observations were made in multiple independent experiments (n=3). (c) Lesion progression in representative leaves upon treatment with the strains mentioned therein at 14 dpi. The yellow horizontal lines mark the leading edge of the disease lesion.

### OsS5H/FNS-03g functionally complements its *Arabidopsis* homologue AtDMR6

Protein sequence analysis revealed that OsS5H/FNS-03g is homologous to an *Arabidopsis thaliana* protein called DOWNY MILDEW RESISTANT 6 (AtDMR6; Figure 3a). AtDMR6 has been characterised as a susceptibility factor for multiple pathogens including *Hyaloperonospora parasitica* (downy mildew; oomycete), *Phytophthora capsica* (an oomycete pathogen), *Pseudomonas syringae* pv. *tomato* DC3000 (*Pst* DC3000; bacterial pathogen) (Zhang et al., 2017). To determine whether OsS5H/FNS-03g shares functional homology with AtDMR6, an *Arabidopsis thaliana* line lacking *AtDMR6* gene (*atdmr6*) was transformed with a construct carrying 35S:*OsS5H/FNS-03g*-*GFP*. Further, the transgenic plants in T_2_ generation, the *atdmr6* mutant, and the wild-type line (Col0) were inoculated with the bacterial pathogen *Pst* DC3000, and bacterial growth was scored subsequently at 2dpi. It was observed that ectopic expression of *OsS5H/FNS-03g* in *atdmr6* restored the susceptibility to *Pst* DC3000 (Figure 3b, c) indicating a functional homology between OsS5H/FNS-03g and AtDMR6 *in planta*.

**Figure 3:**
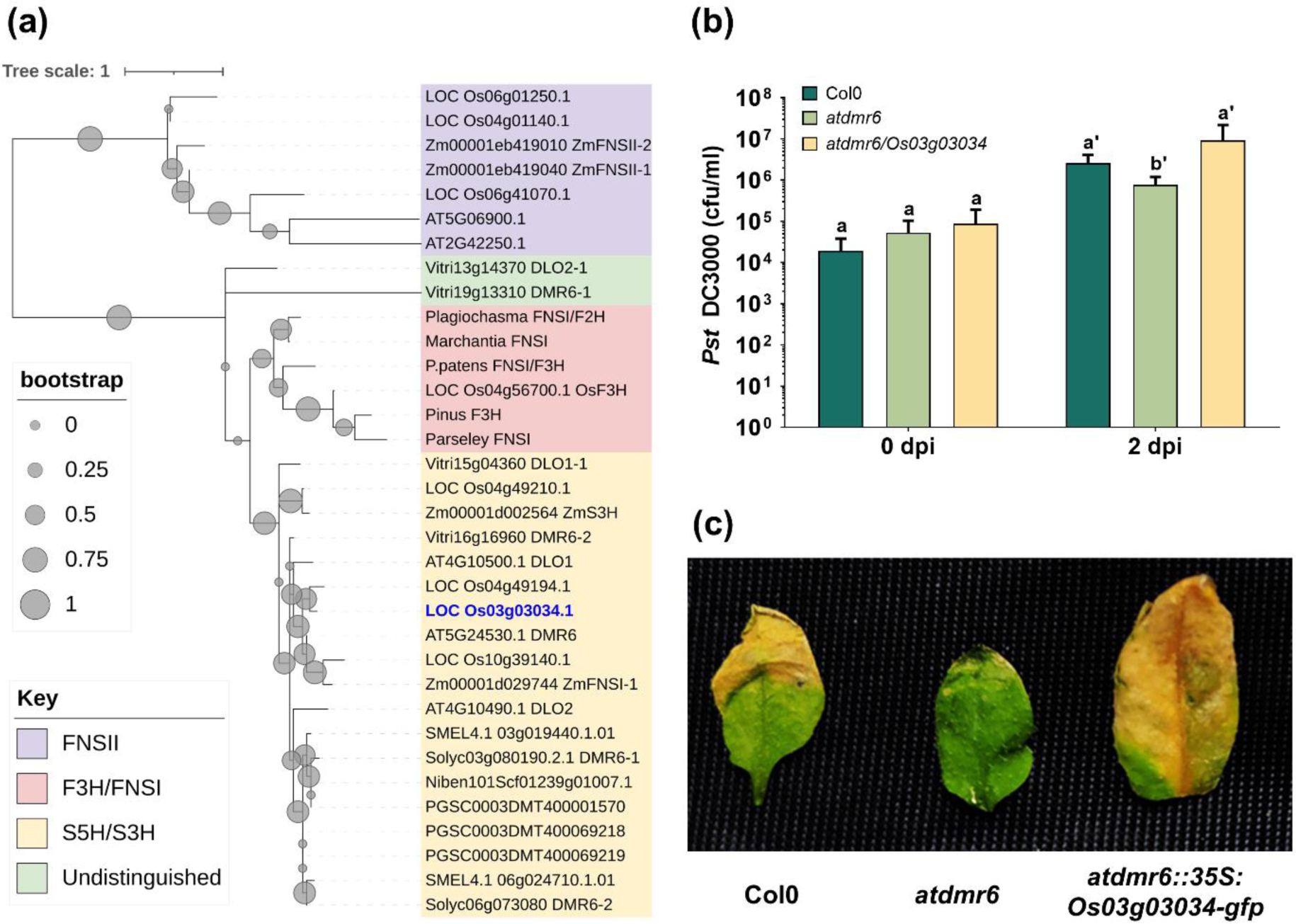
OsS5H/FNS-03g complements AtDMR6 function in *Arabidopsis thaliana*. (a) A neighbourhood-joining based dendrogram indicating the sequence homology between various orthologues of OsS5H/FNS-03g (in blue font). Protein sequences of the indicated species were retrieved from literature information or by protein BLAST using OsS5H/FNS-03g sequence as query. The clades are coloured based on the protein functions (b) Rosette leaves of wildtype *Arabidopsis thaliana* Col0, *atdmr6* mutant, and transgenic *atdmr6* ectopically expressing *OsS5H/FNS-03g-GFP* were syringe infiltrated with *Pst* DC3000 of OD_600nm_=0.02. Bacterial growth after 2 days of infiltration was determined by serial dilution plating on King’s B agar plates. (c) Disease symptoms on the infected leaves at 5 dpi. Bars in (b) represent the mean colony forming units and the error bars indicate standard deviations in at least 3 biological replicates. Bars capped with different alphabets indicate significant differences at a confidence limit of 95% using Student’s *t*-test. Similar results were obtained in three independent experiments.

### OsS5H/FNS-03g is a bifunctional protein with S5H and FNS activities

Complementation analysis confirmed that OsS5H/FNS-03g is a functional homologue of AtDMR6. AtDMR6 is an SA hydroxylase that hydroxylates SA at the 5’ position resulting in the formation of 2,5 dihydroxybenzoic acid (2,5 DHBA) (Zeilmaker et al., 2015). OsS5H/FNS-03g was reported by independent studies to have either S5H or FNS activities (Kim et al., 2008; Liu et al., 2023; Zhang et al., 2022). Therefore, we investigated whether the protein possesses both S5H and FNS activities. To this end, we expressed recombinant OsS5H/FNS-03g that is N-terminally fused to Maltose Binding Protein (MBP) in *Escherichia coli* and purified the protein using amylose resin. The purified protein preparation was used to perform biochemical reactions to test the enzymatic function using SA and Naringenin as substrates in independent reaction conditions. Purified MBP was used as a negative control. Results show that OsS5H/FNS-03g efficiently catalyses SA hydroxylation at the 5’ position thereby producing 2,5-DHBA (Figure 4a). Further, the protein was also found to possess FNS activity which was observed as a peak corresponding to the retention time of Apigenin standard (Figure 4b). To confirm if the actual product is produced, the fraction corresponding to apigenin retention time was collected from the standard, MBP control-, and MBP-OsS5H/FNS-03g-containing reactions to analyse the product using Matrix-Assisted Laser Desorption/Ionisation Time-Of-Flight (MALDI-TOF). The MALDI spectrum revealed a peak with a mass-to-charge (m/z) ratio corresponding to apigenin (m/z = 271; including 1 Da added by positive mode ionization) in standard and MBP-OsS5H/FNS-03g, and not in MBP control sample (Figure S6). The results here suggest that OsS5H/FNS-03g possesses S5H as well as FNS-I activities. Additionally, we noted that SA and Naringenin induce the expression of *OsS5H/FNS-03g* to varied extents (Figure S3b).

**Figure 4:**
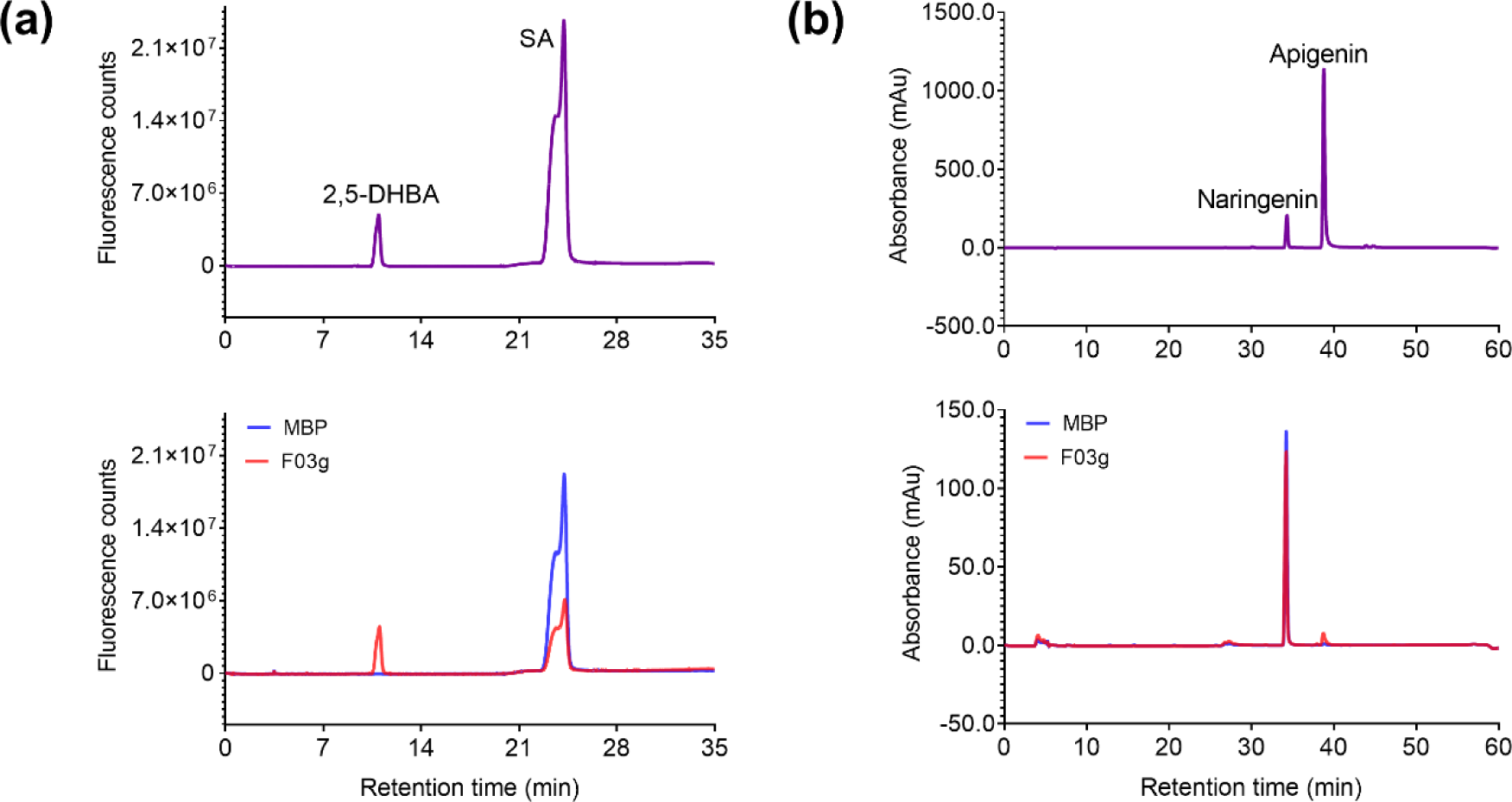
OsS5H/FNS-03g is a bifunctional protein. Chromatograms showing the authentic standards (top) and reaction mixtures of MBP-OsS5H/FNS-03g (F03g) and MBP when incubated with (a) SA and (b) Naringenin. The reactions were performed for one hour at appropriate temperature and reaction conditions. The reaction mixture and authentic standards were resolved further using High Performance Liquid Chromatography to observe the product formation.

### Apigenin promotes virulence of PXO99^A^ *tal9b*-

As OsS5H/FNS-03g catalysed the formation of apigenin, we asked if apigenin plays any role in virulence of Xoo. For this, we exogenously applied 10µM apigenin by spraying on rice leaves and subsequently infected the leaves with PXO99^A^ and PXO99^A^ *tal9b*-(24 hours post spraying). Interestingly, we observed that prior application of apigenin enhanced the virulence of the PXO99^A^ *tal9b*- and did not affect the virulence of PXO99^A^ (Figure 5a, b). These results indicate that apigenin promotes the virulence of an otherwise less virulent mutant of Xoo, which is also less proficient in inducing the expression of *OsS5H/FNS-03g*.

**Figure 5:**
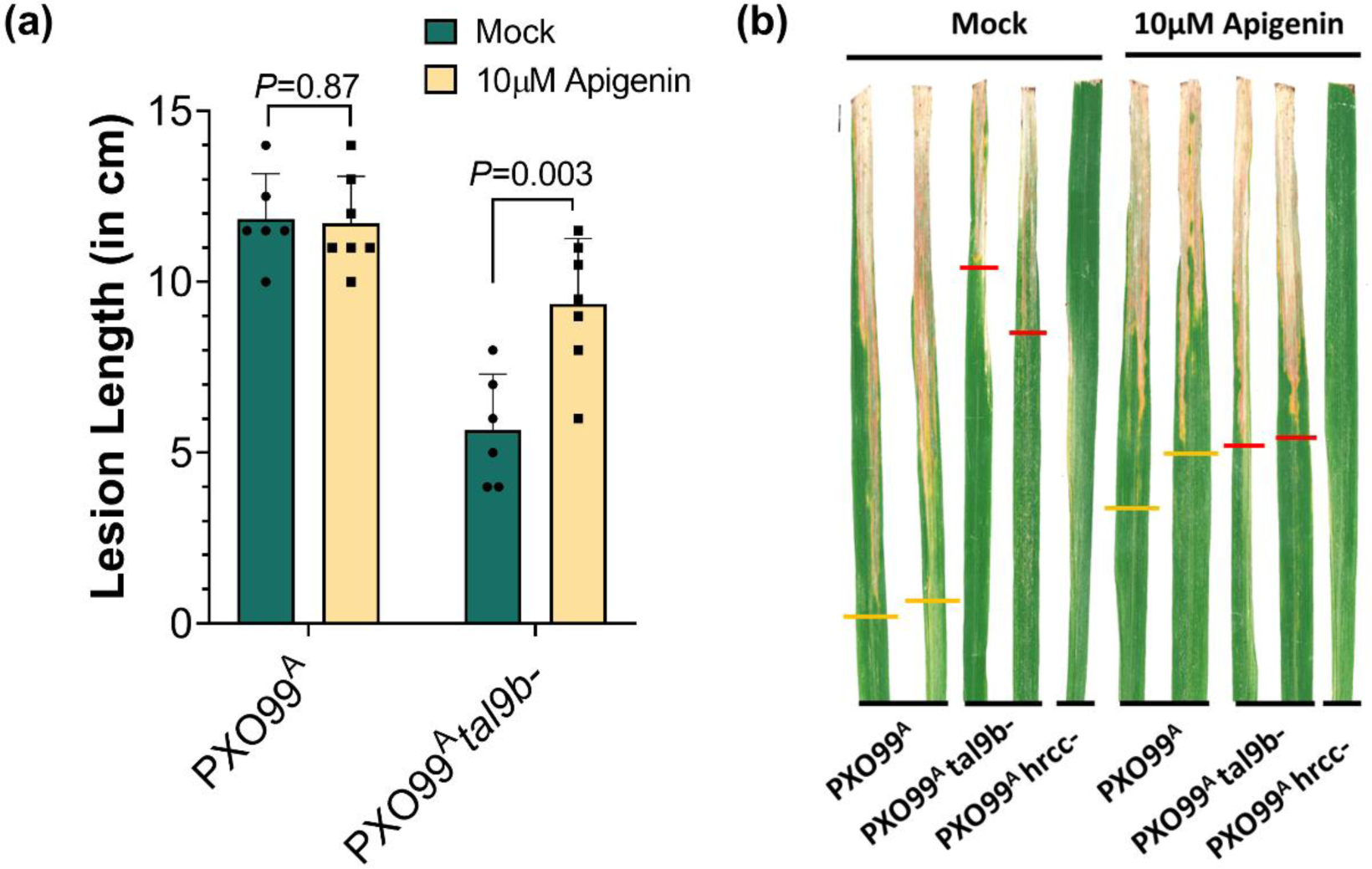
Apigenin promotes virulence of PXO99^A^ *tal9b*-. (a) TN1 rice leaves were sprayed with either mock or 10µM apigenin to run off and were infected with the Xoo strains PXO99^A^ and PXO99^A^ *tal9b*-at 24 hours post spraying. Lesion lengths were measured at 14 dpi and the mean length is represented as bars while the error bars indicate the standard deviation of the lesion length in at least 6 infected leaves from one experimental replicate. The *P-*value was calculated using an unpaired Student’s *t-*test. The experiment was repeated twice with similar observations. (b) The appearance of lesions in the infected leaves at 14 dpi. Yellow and red lines on the leaves mark the tip of the lesion caused by PXO99^A^ and PXO99^A^ *tal9b*-, respectively.

## DISCUSSION

The Xoo pathogen deploys a sophisticated class of type III effector proteins called the TAL effectors to take direct control on the transcription of certain host genes. Several host genes that are activated in a TALE-dependent manner are reported to be *bona fide* susceptibility factors (Boch et al., 2009; Streubel et al., 2013; Tran et al., 2018). Such TALEs are required for full virulence of the pathogen. The Xoo strain PXO99^A^ encodes 19 TALEs in its genome (Salzberg et al., 2008).

Here we report the identification and characterization of *OsS5H/FNS-03g* - a bifunctional protein-coding gene that is activated by the PXO99^A^ TALE TAL9b. We observed that *OsS5H/FNS-03g* is a functional homologue of the *Arabidopsis thaliana* DOWNY MILDEW RESISTANT 6 (AtDMR6) protein. Accumulating evidence indicate that AtDMR6 and its orthologues are disease susceptibility factors for various pathogens in a wide variety of plant species including *A. thaliana*, tomato, grapevine, maize, rice, and sweet basil (Djennane et al., 2023; Falcone Ferreyra et al., 2015; Hasley et al., 2021; Liu et al., 2023; Pirrello et al., 2022; Thomazella et al., 2021; Van Damme et al., 2008; Wu et al., 2022; Zeilmaker et al., 2015; Zhang et al., 2022; Zhu et al., 2020). Biochemically, DMR6 orthologues catabolise Salicylic acid (Thomazella et al., 2021; Zhang et al., 2017). Therefore, it is evident that several evolutionarily distant pathogens with different modes of infection, convergently mediate SA catabolism. This manipulation leads to a reduction in the *in planta* levels of SA, thereby diminishing host immunity and possibly systemic acquired resistance.

### Possible role of flavones in disease susceptibility

OsS5H/FNS-03g was first reported as a Flavone Synthase I (FNSI) that catalyses the formation of the flavone apigenin using the flavanone naringenin as substrate (Kim et al., 2008). As seen in Figure 3, S5H, FNS, and F3H proteins share high sequence homology. In addition, neofunctionalization and dual activities of several F3H homologous proteins have been reported by previous studies in various plant species (Li et al., 2020; Pucker and Iorizzo, 2023). Supporting (Falcone Ferreyra et al., 2015) and contradicting (Thomazella et al., 2021) evidences exist regarding AtDMR6 and its FNS activity. Here, using the same protein preparation, we showed that OsS5H/FNS-03g possesses both S5H and FNS activities *in vitro*. Further, while testing the possible role of apigenin in plant susceptibility to Xoo, we observed that apigenin promoted virulence of PXO99^A^ *tal9b*-, a strain that is otherwise less virulent and defective in inducing *OsS5H/FNS-03g*.

Apigenin is known to serve multiple functions including the protection of plants from Ultraviolet B (UV-B) radiation-induced damages (Righini et al., 2019), enhanced nutritional value (Casas et al., 2014), enhanced tolerance to nitrogen deprivation through microbiome modulation (Yu et al., 2021), enhanced biofilm formation by soil diazotrophic bacteria (Yan et al., 2022), and a putative role in senescence (Falcone Ferreyra et al., 2015). Apigenin is also known as a potent inducer of *nod* genes in *Sinorhizobium meliloti* (Peters et al., 1986; Watson et al., 2015). In a similar manner, apigenin may induce expression of Xoo genes that are involved in interaction with rice. Specifically, as in diazotrophs, apigenin may promote biofilm formation of Xoo and thus enhance virulence. Furthermore, accumulation of apigenin beyond a certain level during infection may also have a role in promoting senescence in leaves and thus, enhance symptom formation. As can be seen, the literature suggests multiple possibilities by which apigenin can promote virulence of Xoo, and additional studies are needed to distinguish these various possibilities.

Yet another functional aspect of Xoo-mediated upregulation of an FNS activity-containing protein could be to reduce the basal levels of the flavanone naringenin. It has been reported that naringenin inhibits the growth of *X. oryzae in vitro* (Padmavati et al., 1997). Further, the accumulation of naringenin was shown to provide broad-spectrum resistance to rice pests (Yang et al., 2021). Additionally, naringenin was found to possess antimicrobial and defense-inducing properties in other plant species like tobacco and *Arabidopsis* (An et al., 2021; Shi et al., 2024; Sun et al., 2022). Therefore, it can be speculated that reducing the basal level of naringenin during pathogenesis, possibly mediated by the induction of *OsS5H/FNS-03g* will aid in rendering the host environment conducive for pathogen growth.

### OsS5H/FNS-03g is convergently targeted by diverse rice pathogens

Xoo TALEs that target *OsS5H/FNS-03g* are present in most of the sequenced strains (Figure S4). Intriguingly, *OsS5H/FNS-03g* is one of the few rice genes that is targeted by TALEs from both the *X. oryzae* pathovars (Xoo and Xoc) by binding to the same effector binding element (Cernadas et al., 2014; Mücke et al., 2019). Mücke et al., (2019) discussed the convergence in the virulence mechanisms of the two *X. oryzae* pathovars and the plant resistance mechanisms at OsS5H/FNS-03g and further called it a hub in the plant-pathogen network. The observation that Xoo and Xoc pathovars are convergently targeting *OsS5H/FNS-03g* denotes the existence of pathogenicity mechanisms that function beyond the tissue specificity of the pathogens. That is, there exist certain susceptibility mechanisms that are central to pathogenesis irrespective of the infection style and tissue specificity of the infecting pathogen. This is further substantiated by the observation that the expression of *OsS5H/FNS-03g* is induced by rice pathogens from different kingdoms (Figure S2). Identifying and characterizing such broad-spectrum disease susceptibility factors will inform us of targets for highly durable disease resistance breeding.

Overall, our study suggests that the flavone apigenin might play some important roles in determining the outcomes of plant-pathogen interaction. Future studies directed towards understanding the *in planta* and in bacteria role of apigenin are hence required. Also, the disruption of EBE in the upstream sequence of *OsS5H/FNS-03g* through genome editing approaches and pyramiding the disrupted EBE with other widely used resistance alleles can be explored for developing broad spectrum resistance to Xoo.

## EXPERIMENTAL PROCEDURES

### Plant, bacterial materials, and growth conditions

The rice variety Taichung Native 1 (TN1), which is susceptible to bacterial blight disease, was used for gene expression analysis and Xoo virulence assays. The seeds were washed, soaked, and germinated on a moist filter paper and grown on Petri plates for 7 days. Later, the seedlings were transplanted to pots containing black soil and grown for 7 more days for gene expression analysis or were transplanted to the field until they were 60-days old for performing virulence assays. All the experiments concerning TN1 were performed in green house conditions with temperature not exceeding 30°C with 12 hour:12 hour light: dark cycles and natural light conditions. *Arabidopsis thaliana* wildtype accession Columbia-0 (Col-0) and the T-DNA insertion mutant line *atdmr6* (NASC ID: SK19087) were germinated on one-strength Murashige-Skoog Agar medium with 2% sucrose and 0.8% agar at 22°C/18°C and 16h light/8h dark cycles. *atdmr6* lines were selected on plates containing an appropriate amount of phosphinothricin. *Arabidopsis* seedlings were grown in a plant growth chamber (AR-75L3, Percival-Scientific, USA) at 22°C/18°C at 16h light/8h dark cycles in pots containing 1:1:1 ratio of perlite: vermiculite: soilrite C (Keltech Energies, India).

Bacterial strains used in this study include *Escherichia coli* DH5α, S17-1 and Rosetta DE3, *Agrobacterium tumefaciens* AGL1, *Xanthomonas oryzae* pv. *oryzae* PXO99^A^, PXO86 and BXO43, *X. o*. pv. *oryzicola* BLS256 and BXORI, and *Pseudomonas syringae* pv. *tomato* DC3000. *E. coli* strains were grown on Luria Bertani medium (10g/L peptone, 5g/l yeast extract, 10g/L sodium chloride, pH 7.0); *Agrobacterium* strains were cultured on Yeast Extract Mannitol medium (1g/L yeast extract, 10g/L Mannitol, 0.5g/L dipotassium phosphate, 0.2g/L magnesium sulphate, 0.1g/L sodium chloride, pH 7.0); Xoo and Xoc were cultured on PS medium (1% peptone and 1% sucrose, pH 7.0); Pst DC3000 was cultured in King’s B medium. E. coli was cultured at 37°C while the other bacteria were grown at 28°C. Appropriate antibiotics/selection markers were used at the following concentrations: Rifampicin 50µg/ml, Kanamycin 50µg/ml, Carbenicillin 50µg/ml, Chloramphenicol 30µg/ml, Spectinomycin 50µg/ml, Phosphinothricin (Basta) 20µg/ml, Hygromycin 25µg/ml. All the bacterial strains, plant materials and plasmids used in this study are listed in Table S1.

### Analysis of public transcriptome data

Publicly available Affymetrix microarray data pertinent to the interaction between rice and its biotic stress factors with appropriate controls were retrieved from the NCBI-Gene Expression Omnibus (GEO) repository (Table S2). The data were normalised and analysed using Affymetrix Expression Console and Transcriptome Analysis Console. All the datasets were normalised using the PLIER algorithm and further analysed to identify the differentially expressed genes (|FC|≥1.5 and *P-*value≤0.05). The fold change values of *LOC_Os03g03034* were retrieved by searching for the probe ID of the gene: Os.10510.1.S1_at and plotted in R-studio using the *ggplot2* library. The expression level (FPKM) of *LOC_Os03g03034* under various biotic stresses was retrieved from the Rice RNA-seq database [(http://ipf.sustech.edu.cn/pub/ricerna/; (Zhang et al., 2020)] and further processed and plotted using Microsoft Excel and GraphPad Prism 8, respectively.

### Generation of mutant strains of Xoo and Xoc

To generate the type-3 secretion system deficient strains of Xoo and Xoc, a 506bp long segment of the *hrcC* gene was amplified from the genomic DNA of Xoo PXO99^A^ and Xoc BXORI and cloned between the EcoRI and BamHI sites of the mobilizable vector pK18mob, which upon integration into the bacterial genome, disrupts the gene and confers the cells Kanamycin resistance. The plasmid DNA were sequence confirmed and used to transform the *E. coli* strain S17-1. Further a biparental mating was set between *E. coli* S17-1 carrying the plasmid of interest and the Xoo/Xoc strains at a ratio of 1:5, 1:15, and 1:25 of *E. coli* to Xoo/Xoc for 48-72 hours on autoclaved nylon membranes at 28°C. The cells were scraped and plated on PS plates containing Kanamycin to select the integrants. The integration was confirmed through PCR with appropriate primer pairs. Generation of *Tal9b* mutant of PXO99^A^ was generated in a similar manner using a 1252 bp fragment encoding a part of the DNA-binding domain of the TALE for homology-based marker integration. All the primers used in this study are listed in Table S3.

### Gene expression analysis

Xoo and Xoc strains (wildtype and mutants) were cultured in PS broth for 24 hours at 28°C. The cells were pelleted by centrifugation at 4000xg for 10 minutes, washed with water and adjusted to an optical density of 1.0 at 600nm with autoclaved water. The cell suspensions or autoclaved water (mock) were syringe-infiltrated into 14 days-old TN1 leaves and further incubated in the green house. Leaves around the site of infiltration (2cm) were collected 24 hours post infiltration, flash-frozen in liquid Nitrogen, and stored at −80°C until further processing. For clip-inoculations, bacterial suspensions were processed as mentioned above and fully expanded leaves of TN1 plants (50-60 days old) were cut with a scissor dipped in the bacterial suspension. Leaf pieces of 1cm length were collected from the site of clipping at different time points and stored at −80°C until further processing. RNA was isolated from the leaves (8-10 leaves per treatment for infiltrated samples; 4-5 leaf pieces from clip-inoculated samples) using RNeasy Plant Mini Kit (Qiagen, Germany). Depending on the RNA concentration, about 1µg to 5µg total RNA was used for first-strand synthesis using RNA to cDNA EcoDry Premix with Oligo(dT) (Takara Bio Inc., Japan), by following the manufacturer’s recommendations. cDNA was diluted to an RNA-equivalent concentration of 20-25ng/µl using nuclease-free water. Quantitative real-time PCR (qRT-PCR) reactions were performed using 1ul diluted cDNA as template and 2X *Power* SYBR™ Green PCR Master Mix (Applied Biosystems, US) in a 10µl reaction with 0.5µM forward and reverse primers each. The reactions were performed on a ViiA 7 Real-Time PCR System (Applied Biosystems, US) or CFX384 Touch RT-PCR detection system (Bio-Rad, US). *OsGAPDH* (*LOC_Os04g40950.1*) was used as the internal control for expression normalisation. Relative fold change values were calculated using the 2^-ΔΔCT^ method (Livak and Schmittgen, 2001). For semi-quantitative PCR, a 20µl reaction containing 2X KAPA Taq ReadyMix (Roche Applied Sciences, Germany), 0.5µM forward and reverse primers specific to the transcript variants/*OsGAPDH* and 1µl cDNA template (20ng/µl RNA-equivalent) was set and cycled 20 or 25 times. The reactions were then resolved in a 2% agarose gel and documented.

### Xoo virulence assay

Xoo strains were cultured and processed as described above and the cell suspensions were used to inoculate fully expanded leaves of 60-days old TN1 rice plants using the clip-inoculation method with a scissor dipped in bacterial suspension (Kauffman et al., 1973). Lesion lengths were measured at 14 days post inoculation from the site of clipping to the leading edge of the disease lesion (dpi).

### *in silico* analyses

The EBE sequence and its corresponding TAL effector was predicted using the daTALbase server (Pérez-Quintero et al., 2018). The RVD sequences and classes of all the Xoo and Xoc TAL effectors were retrieved using the “*Load and View TALE classes*” function of the AnnoTALE tool (Grau et al., 2016). For the sequence homology analysis of LOC_Os03g03034.1, all the known DMR6 homologues were identified and retrieved from corresponding publications. The FNS, F2H, and F3H sequences were obtained from Li et al, 2020. Where required, the homologous sequences were obtained using NCBI BLASTp function with LOC_Os03g03034.1 sequence as query. Phylogenetic analysis of the protein sequences was conducted using the *Phylogeny.fr* server using default settings (Dereeper et al., 2008) and the tree was visualised and annotated using the iTOL server (Letunic and Bork, 2021). All the sequences used for the phylogenetic analysis are provided in Data S1.

### Cloning *OsS5H/FNS-03g* for plant and biochemical experiments

The coding sequence of *LOC_Os03g03034.1* was amplified from rice cDNA and cloned into the Gateway-compatible entry vector pENTR/D-TOPO (Invitrogen, US) and sequence verified. Further, the gene was sub-cloned into the Gateway destination vector pH7FWG2 using LR Clonase II enzyme mix (Invitrogen, US) to generate the overexpression (CaMV 35S promoter) expression clone with C-terminally fused eGFP. For recombinant protein expression, the CDS of *LOC_Os03g03034.1* was cloned into pETM-40 using restriction-free cloning (van den Ent and Löwe, 2006) to generate a reading frame comprising *Maltose Binding Protein (MBP)*, *LOC_Os03g03034.1*, and *6X Histidine* coding sequences.

### Generation of *Arabidopsis* transgenics and *Pseudomonas syringae* infection assay

*A. tumefaciens* AGL1 carrying the pH7FWG2:*Os03g03034.1* construct was cultured overnight with appropriate antibiotics. An overnight grown, 250ml *Agrobacterial* culture was centrifuged at 4000xg for 10 minutes, washed with water, and resuspended in 250ml floral-dip solution (½ strength MS, 5% sucrose, 0.01% Silwet L-77). Developing inflorescence of *Arabidopsis thaliana atdmr6* plants were dipped into the bacterial suspension for 2 minutes and kept under high humidity for 24 hours and were maintained until seed collection at 22°C and 16-hour photoperiod (Clough and Bent, 1998). The collected T_0_ seeds were surface sterilised for 20 minutes in 70% ethyl alcohol, stratified at 4°C for 48 hours, and plated on MS agar plates containing appropriate selection (Phosphinothricin and Hygromycin). The selected plants were further transferred to soil and grown for genotyping and collecting T_1_ seeds. The collected seeds were grown on selection plates for further experiments. The primers used for genotyping the mutant plants and transgenics are listed in Table S3. Expression of Os03g03034-GFP in the transgenic leaves was confirmed through immunoblotting using an anti-GFP antibody (Abcam ab6556; 1:3000 dilution) and HRP-conjugated anti-rabbit secondary antibody (Sigma 12-348; 1:50,000 dilution) (Figure S7). Freshly grown Pst DC3000 cells were resuspended in autoclaved water and adjusted to an OD_600_ of 0.02. The cell suspension was syringe-infiltrated into the fully expanded rosette leaves of Col-0, *atdmr6*, and *atdmr6/Os03g03034-gfp* and maintained at 22°C and 16-hour photoperiod under high humidity. The bacterial population was quantified at 0 dpi and 2 dpi by serial dilution plating and counting the number of colony forming units per ml (cfu/ml) leaf extract.

### Recombinant protein purification

The plasmids pETM-40 and pETM-40:*Os03g03034.1* were mobilised into *E. coli* Rosetta DE3 cells. Both the clones were cultured at 37°C to an optical density of 0.4-0.6 at 600nm and the protein expression was induced by adding 0.5mM Isopropyl ß-D-1-thiogalactopyranoside (IPTG). The induced cultures were further grown at 18°C and 200rpm for 16-20 hours. The proteins were purified using the amylose resin affinity purification method by following the manufacturer’s recommendations (E8021L, NEB). Briefly, about 1.5mL slurry was used for purifying proteins from a 250mL culture. The induced cells were pelleted at 4000xg for 20 minutes and the pellet was resuspended in 10mL lysis buffer (20mM Tris-HCl (pH 7.4), 1mM EDTA, 200mM NaCl, 1mM PMSF, 1X protease inhibitor cocktail, 1mg/ml Lysozyme, 1mM dithiothreitol). The cell suspension was sonicated at 30% amplitude for 20-30 minutes with 10 sec on-off cycles. The lysate was clarified by centrifugation at max speed for ∼30 minutes. The supernatant was mixed with equilibrated resin and was gently mixed for 3 hours at 4°C and10rpm in a rotator. The unbound fraction was collected by spinning the tubes at 100xg for 1-2 minutes. The beads were washed with 5 bead volumes of wash buffer (20mM Tris-HCl (pH 7.4), 1mM EDTA, 200mM NaCl) five times. The protein elution was performed with 2 ml elution buffer (wash buffer+10mM maltose) each 3 to 4 times. Washing and elution were performed by incubating the beads in corresponding buffers for 5-10 minutes at 4°C and RT, respectively. All the fractions were resolved on an SDS-PAGE to assess protein recovery and purity. Protein samples were dialyzed against dialysis buffer (50mM Tris (pH 7.4), 100mM KCl, 1mM DTT, 30% Glycerol) 7 times and concentrated to a volume of ∼600ul using Amicon Unltra-155 centrifugal filters with MW cut-off of 30kDa. Protein quantity was measured using NanoDrop Lite (Thermo Scientific, US). The concentrated samples were stored at −30°C until further experiments.

### Biochemical assay for S5H activity

The biochemical assay to test the SA hydroxylation activity of the protein was performed as described earlier (Thomazella et al., 2021). Briefly, 60µg purified protein (MBP and MBP-OsS5H/FNS-03g) was incubated in a 200µl reaction mixture at 40°C for 1-hour (50mM 2-(N-morpholino) ethanesulfonic acid (MES; pH 6.5), 1mM α-ketoglutarate, 10mM ascorbic acid, 0.4mM ferrous sulphate, 7.2µM salicylic acid) and later filtered through 0.22µm filters. The filtered reaction mixture (10µl) was passed through a Zorbax SB C18 reverse phase column (4.6mm x 25cm), 5um pore size at a flow rate of 0.75ml/min for a run time of 35 minutes per sample in a Thermo Vanquish UHPLC system. The column was first equilibrated with 100% methanol. The mobile phase used for the run includes A: 0.2M sodium acetate (pH 5.5) and B: 100% methanol. The run condition is as follows: A:97%-B:3% for 15 minutes, followed by A:93%-B:7% for 15 minutes, followed by A:97%-B:3% for 5 minutes. Authentic standards of SA (#247588; Sigma, USA) and 2,5-DHBA (#149357; Sigma, USA) were dissolved in 100% dimethyl formamide (DMF) at the required concentration (∼7.2µM). SA was detected at 403nm, while 2,5-DHBA was detected at 438nm, using a fluorescence detector. The retention time of SA was observed to be ∼24 minutes and that of 2,5-DHBA was ∼11 minutes.

### Biochemical assay for FNS activity

The biochemical assay to test the FNS activity of the protein was performed as follows: 100µg purified protein (MBP and MBP-OsS5H/FNS-03g) was incubated in a 200µl reaction mixture at 30°C for 1 hour (50mM Tris (pH 7), 160µM α-ketoglutarate, 1mM ascorbic acid, 100µM ferrous sulphate, 200µM naringenin) and filtered through 0.22µm filters. The filtered reaction mixture (50µl) was resolved on a Thermo Dionex UltiMate 3000 system using a Zorbax SB C18 reverse phase column (4.6mm x 25cm), 5um pore size at a flow rate of 0.5ml/min for a run time of 1 hour per sample. The column was first equilibrated with 100% methanol. The mobile phase used for the run includes A: 80% 10mM Ammonium acetate (pH 5.6): 20% Methanol and B: 100% methanol. The flow was set to 0% B for 2 minutes and 0 to 100% B in 50 minutes followed by 100% to 0% B in 3 minutes and continued at B 0% for 5 minutes. Authentic standards of Naringenin (#N5893; Sigma, USA) and Apigenin (#10798; Sigma, USA) were dissolved in 100% DMF at the required concentration (∼27µM). Naringenin and apigenin were detected at 292nm and 340nm, respectively, using a UV detector. The retention time of naringenin was observed to be ∼33 minutes and that of apigenin was ∼38 minutes.

Fractions corresponding to naringenin and apigenin retention times were collected using a Thermo Scientific Fraction Collector and vacuum dried using Concentrator Plus (Eppendorf, Germany). Dried fractions were resuspended in x ul of 70% acetonitrile with 0.1% trifluoroacetic acid. 1 µl of the sample and 1 µl of 10 mg/ml CHCA (α-Cyano-hydroxycinnamic acid) matrix in 70% acetonitrile with 0.1% trifluoroacetic acid were mixed and spotted on a MALDI plate and air-dried. The electrospray ionization (ESI) was performed in positive ion mode. Mass spectra were acquired over a mass range of m/z 200-400. MALDI-TOF spectra were acquired using a Spiral TOF JMS-S3000 (JEOL, Japan) Mass Spectrometer.

### Chemical treatment to assess Xoo virulence and gene expression levels

Fully expanded leaves of 60 days-old TN1 plants were sprayed with 10µM apigenin (in water and 0.05% Tween-20) or autoclaved water alone (with the appropriate amount of DMF and 0.05% Tween-20) until run-off and grown in a green-house for 24 hours. Later, the leaves were clip inoculated with the wild type or mutant strains of Xoo using the clip-inoculation method and the lesion length was measured as described earlier.

Treatment with SA and naringenin was performed by incubating detached leaf pieces (1cm^2^) from 50-60 days old TN1 plants on solutions containing 1mM SA or 0.5mM naringenin or solvent control (mock). The stock solutions were prepared in absolute methanol and diluted to the required concentration in autoclaved Milli-Q water with 0.01% Silwet^TM^ L-77 (Momentive, NY, USA). The leaf pieces were collected at 1 hour, 6 hours, 12 hours, and 24 hours post-treatment, flash frozen, and stored at −80°C until further processing.

### Statistical analysis

The data generated in this study were analysed using appropriate statistical methods, wherever necessary, and the details are provided in the corresponding figure legends.

### Accession numbers

OsS5H/FNS-03g - LOC_Os03g03034; OsGAPDH - LOC_Os04g40950; OsSWEET11 - LOC_Os08g42350; OsTFX1 - LOC_Os09g29820; hrcC (PXO99^A^) - PXO_RS00410; Tal9b (PXO99^A^) - PXO_RS02485; hrcC (BXOR1) - ACU15_01175; AtDMR6 - AT5G24530. *atdmr6* mutant plant *-* SK19087.

## ACKNOWLEDGEMENTS

This study was supported by the grants provided to HKP from the Council of Scientific and Industrial Research, New Delhi, India (MLP0121 Phase-I and II) and to RVS from the Science and Engineering Research Board (SERB), Department of Science and Technology, Government of India (SB/S2/JCB-12/2014). GCG, SD, NG, AM, NG, RPR, and NS thank the Council for Scientific and Industrial Research for their fellowships. DJ thanks the Department of Biotechnology, New Delhi, India for granting their Research Associateship.

## DATA AVAILABILITY STATEMENT

Information on publicly available data analysed in this study is provided in the supplementary information of this article. All other data generated for this study are present in the main text or supplementary information. The materials from this study can be obtained from the corresponding author upon reasonable request.

## AUTHOR CONTRIBUTION

Conceptualisation - GCG, SD, and HKP. Investigation - GCG (computational and phylogenetic analyses), GCG, SD, NG (bacterial mutagenesis, gene expression analysis, gene cloning for plant expression, *Arabidopsis* complementation, pathogen infection assays); GCG, AM, GN (recombinant protein purification); AM (biochemical activity assays, HPLC, and gene expression analysis); NS (gene cloning for protein purification); RPR and DJ (plant transformation). Assessment and supervision of the work - HKP and RVS. Writing of original draft – GCG, SD, NG. Manuscript proofreading - all the authors. Funding acquisition – HKP and RVS.

## REFERENCES

An, J., Kim, Sun Ho, Bahk, S., Vuong, U.T., Nguyen, N.T., Do, H.L., et al. (2021) Naringenin Induces Pathogen Resistance Against Pseudomonas syringae Through the Activation of NPR1 in Arabidopsis. Frontiers in Plant Science, 12, 672552. 10.3389/fpls.2021.672552.

Antony, G., Zhou, J., Huang, S., Li, T., Liu, B., White, F., et al. (2010) Rice xa13 recessive resistance to bacterial blight is defeated by induction of the disease susceptibility gene Os-11N3. Plant Cell, 22, 3864–3876. 10.1105/tpc.110.078964.

Bezrutczyk, M., Yang, J., Eom, J.-S., Prior, M., Sosso, D., Hartwig, T., et al. (2018) Sugar flux and signaling in plant-microbe interactions. The Plant Journal: For Cell and Molecular Biology, 93, 675–685. 10.1111/tpj.13775.

Blanvillain-Baufumé, S., Reschke, M., Solé, M., Auguy, F., Doucoure, H., Szurek, B., et al. (2017) Targeted promoter editing for rice resistance to Xanthomonas oryzae pv. oryzae reveals differential activities for SWEET14-inducing TAL effectors. Plant Biotechnology Journal, 15, 306–317. 10.1111/pbi.12613.

Boch, J., Scholze, H., Schornack, S., Landgraf, A., Hahn, S., Kay, S., et al. (2009) Breaking the code of DNA binding specificity of TAL-type III effectors. Science, 326, 1509– 1512. 10.1126/science.1178811.

Bogdanove, A.J., Schornack, S. & Lahaye, T. (2010) TAL effectors: Finding plant genes for disease and defense. Current Opinion in Plant Biology, 13, 394–401. 10.1016/j.pbi.2010.04.010.

Cai, L., Cao, Y., Xu, Z., Ma, W., Zakria, M., Zou, L., et al. (2017) A Transcription Activator-Like Effector Tal7 of Xanthomonas oryzae pv. Oryzicola Activates Rice Gene Os09g29100 to Suppress Rice Immunity. Scientific Reports, 7, 1–13. 10.1038/s41598-017-04800-8.

Casas, M.I., Duarte, S., Doseff, A.I. & Grotewold, E. (2014) Flavone-rich maize: an opportunity to improve the nutritional value of an important commodity crop. Frontiers in Plant Science, 5.

Cernadas, R.A., Doyle, E.L., Niño-Liu, D.O., Wilkins, K.E., Bancroft, T., Wang, L., et al. (2014) Code-assisted discovery of TAL effector targets in bacterial leaf streak of rice reveals contrast with bacterial blight and a novel susceptibility gene. PLoS pathogens, 10, e1003972. 10.1371/journal.ppat.1003972.

Chu, Z., Fu, B., Yang, H., Xu, C., Li, Z., Sanchez, A., et al. (2006) Targeting xa13, a recessive gene for bacterial blight resistance in rice. Theoretical and Applied Genetics, 112, 455–461. 10.1007/s00122-005-0145-6.

Clough, S.J. & Bent, A.F. (1998) Floral dip: a simplified method for Agrobacterium-mediated transformation of Arabidopsis thaliana. The Plant Journal: For Cell and Molecular Biology, 16, 735–743. 10.1046/j.1365-313x.1998.00343.x.

Dereeper, A., Guignon, V., Blanc, G., Audic, S., Buffet, S., Chevenet, F., et al. (2008) Phylogeny.fr: robust phylogenetic analysis for the non-specialist. Nucleic Acids Research, 36, W465–469. 10.1093/nar/gkn180.

Djennane, S., Gersch, S., Le-Bohec, F., Piron, M.-C., Baltenweck, R., Lemaire, O., et al. (2023) Reduced susceptibility to downy mildew in grapevine plants edited for VvDMR6-1. *Journal of Experimental Botany*, erad487. 10.1093/jxb/erad487.

Ent, F. van den & Löwe, J. (2006) RF cloning: a restriction-free method for inserting target genes into plasmids. Journal of Biochemical and Biophysical Methods, 67, 67–74. 10.1016/j.jbbm.2005.12.008.

Falcone Ferreyra, M.L., Emiliani, J., Rodriguez, E.J., Campos-Bermudez, V.A., Grotewold, E. & Casati, P. (2015) The Identification of Maize and Arabidopsis Type I FLAVONE SYNTHASEs Links Flavones with Hormones and Biotic Interactions. Plant Physiology, 169, 1090–1107. 10.1104/pp.15.00515.

Fukagawa, N.K. & Ziska, L.H. (2019) Rice: Importance for Global Nutrition. Journal of Nutritional Science and Vitaminology, 65, S2–S3. 10.3177/jnsv.65.S2.

Grau, J., Reschke, M., Erkes, A., Streubel, J., Morgan, R.D., Wilson, G.G., et al. (2016) AnnoTALE: bioinformatics tools for identification, annotation and nomenclature of TALEs from Xanthomonas genomic sequences. Scientific Reports, 6, 21077. 10.1038/srep21077.

Hasley, J.A.R., Navet, N. & Tian, M. (2021) CRISPR/Cas9-mediated mutagenesis of sweet basil candidate susceptibility gene ObDMR6 enhances downy mildew resistance Varshney, G. (Ed.). PLOS ONE, 16, e0253245. 10.1371/journal.pone.0253245.

Kauffman, H.E., Reddy, A.P.K., Hsieh, S.P.Y. & Merca, S.D. (1973) An improved technique for evaluating resistance of rice varieties to Xanthomonas oryzae. Plant Dis. Rep, 57, 737–741.

Kim, J.H., Cheon, Y.M., Kim, B.-G. & Ahn, J.-H. (2008) Analysis of flavonoids and characterization of theOsFNS gene involved in flavone biosynthesis in Rice. Journal of Plant Biology, 51, 97–101. 10.1007/BF03030717.

Kumar, S., Stecher, G., Li, M., Knyaz, C. & Tamura, K. (2018) MEGA X: Molecular Evolutionary Genetics Analysis across Computing Platforms. Molecular Biology and Evolution, 35, 1547–1549. 10.1093/molbev/msy096.

Letunic, I. & Bork, P. (2021) Interactive Tree Of Life (iTOL) v5: an online tool for phylogenetic tree display and annotation. Nucleic Acids Research, 49, W293–W296. 10.1093/NAR/GKAB301.

Li, D.-D., Ni, R., Wang, P.-P., Zhang, X.-S., Wang, P.-Y., Zhu, T.-T., et al. (2020) Molecular Basis for Chemical Evolution of Flavones to Flavonols and Anthocyanins in Land Plants. Plant Physiology, 184, 1731–1743. 10.1104/pp.20.01185.

Liu, X., Yu, Y., Yao, W., Yin, Z., Wang, Y., Huang, Z., et al. (2023) CRISPR/Cas9-mediated simultaneous mutation of three salicylic acid 5-hydroxylase (OsS5H) genes confers broad-spectrum disease resistance in rice. Plant Biotechnology Journal, 21, 1873–1886. 10.1111/pbi.14099.

Livak, K.J. & Schmittgen, T.D. (2001) Analysis of relative gene expression data using real-time quantitative PCR and the 2(-Delta Delta C(T)) Method. Methods (San Diego, Calif.), 25, 402–408. 10.1006/meth.2001.1262.

Mansueto, L., Fuentes, R.R., Borja, F.N., Detras, J., Abrio-Santos, J.M., Chebotarov, D., et al. (2017) Rice SNP-seek database update: new SNPs, indels, and queries. Nucleic Acids Research, 45, D1075–D1081. 10.1093/NAR/GKW1135.

Mücke, S., Reschke, M., Erkes, A., Schwietzer, C.-A., Becker, S., Streubel, J., et al. (2019) Transcriptional Reprogramming of Rice Cells by Xanthomonas oryzae TALEs. Frontiers in Plant Science, 10.

Niño-Liu, D.O., Ronald, P.C. & Bogdanove, A.J. (2006) Xanthomonas oryzae pathovars: model pathogens of a model crop. Molecular Plant Pathology, 7, 303–324. 10.1111/J.1364-3703.2006.00344.X.

Oliva, R., Ji, C., Atienza-Grande, G., Huguet-Tapia, J.C., Perez-Quintero, A., Li, T., et al. (2019) Broad-spectrum resistance to bacterial blight in rice using genome editing. Nature Biotechnology, 37, 1344–1350. 10.1038/s41587-019-0267-z.

Padmavati, M., Sakthivel, N., Thara, K.V. & Reddy, A.R. (1997) Differential sensitivity of rice pathogens to growth inhibition by flavonoids. Phytochemistry, 46, 499–502. 10.1016/S0031-9422(97)00325-7.

Peng, Z., Hu, Y., Zhang, J., Huguet-Tapia, J.C., Block, A.K., Park, S., et al. (2019) Xanthomonas translucens commandeers the host rate-limiting step in ABA biosynthesis for disease susceptibility. Proceedings of the National Academy of Sciences, 116, 20938–20946. 10.1073/pnas.1911660116.

Pérez-Quintero, A.L., Lamy, L., Zarate, C.A., Cunnac, S., Doyle, E., Bogdanove, A., et al. (2018) daTALbase: A Database for Genomic and Transcriptomic Data Related to TAL Effectors. Molecular Plant-Microbe Interactions®, 31, 471–480. 10.1094/MPMI-06-17-0153-FI.

Peters, N.K., Frost, J.W. & Long, S.R. (1986) A Plant Flavone, Luteolin, Induces Expression of *Rhizobium meliloti* Nodulation Genes. Science, 233, 977–980. 10.1126/science.3738520.

Pirrello, C., Malacarne, G., Moretto, M., Lenzi, L., Perazzolli, M., Zeilmaker, T., et al. (2022) Grapevine DMR6-1 Is a Candidate Gene for Susceptibility to Downy Mildew. Biomolecules, 12, 182. 10.3390/biom12020182.

Pucker, B. & Iorizzo, M. (2023) Apiaceae FNS I originated from F3H through tandem gene duplication. PloS One, 18, e0280155. 10.1371/journal.pone.0280155.

Righini, S., Rodriguez, E.J., Berosich, C., Grotewold, E., Casati, P. & Falcone Ferreyra, M.L. (2019) Apigenin produced by maize flavone synthase I and II protects plants against UV-B-induced damage. Plant, Cell & Environment, 42, 495–508. 10.1111/pce.13428.

Salzberg, S.L., Sommer, D.D., Schatz, M.C., Phillippy, A.M., Rabinowicz, P.D., Tsuge, S., et al. (2008) Genome sequence and rapid evolution of the rice pathogen Xanthomonas oryzae pv. oryzae PXO99A. BMC Genomics, 9, 204. 10.1186/1471-2164-9-204.

Savary, S., Willocquet, L., Pethybridge, S.J., Esker, P., McRoberts, N. & Nelson, A. (2019) The global burden of pathogens and pests on major food crops. Nature Ecology and Evolution, 3, 430–439. 10.1038/s41559-018-0793-y.

Shi, H., Jiang, J., Yu, W., Cheng, Y., Wu, S., Zong, H., et al. (2024) Naringenin restricts the colonization and growth of Ralstonia solanacearum in tobacco mutant KCB-1. *Plant Physiology*, kiae185. 10.1093/plphys/kiae185.

Streubel, J., Pesce, C., Hutin, M., Koebnik, R., Boch, J. & Szurek, B. (2013) Five phylogenetically close rice SWEET genes confer TAL effector-mediated susceptibility to Xanthomonas oryzae pv. oryzae. The New Phytologist, 200, 808–819. 10.1111/nph.12411.

Sugio, A., Yang, B., Zhu, T. & White, F.F. (2007) Two type III effector genes of *Xanthomonas oryzae* pv. *oryzae* control the induction of the host genes *OsTFIIA* γ *1* and *OsTFX1* during bacterial blight of rice. Proceedings of the National Academy of Sciences, 104, 10720–10725. 10.1073/pnas.0701742104.

Sun, M., Li, L., Wang, C., Wang, L., Lu, D., Shen, D., et al. (2022) Naringenin confers defence against Phytophthora nicotianae through antimicrobial activity and induction of pathogen resistance in tobacco. Molecular Plant Pathology, 23, 1737–1750. 10.1111/mpp.13255.

Thomazella, D.P. de T., Seong, K., Mackelprang, R., Dahlbeck, D., Geng, Y., Gill, U.S., et al. (2021) Loss of function of a DMR6 ortholog in tomato confers broad-spectrum disease resistance. Proceedings of the National Academy of Sciences of the United States of America, 118, e2026152118. 10.1073/pnas.2026152118.

Tran, T.T., Pérez-Quintero, A.L., Wonni, I., Carpenter, S.C.D., Yu, Y., Wang, L., et al. (2018) Functional analysis of African Xanthomonas oryzae pv. oryzae TALomes reveals a new susceptibility gene in bacterial leaf blight of rice. PLoS pathogens, 14, e1007092. 10.1371/journal.ppat.1007092.

Van Damme, M., Huibers, R.P., Elberse, J. & Van Den Ackerveken, G. (2008) Arabidopsis *DMR6* encodes a putative 2OG-Fe(II) oxygenase that is defense-associated but required for susceptibility to downy mildew. The Plant Journal, 54, 785–793. 10.1111/j.1365-313X.2008.03427.x.

Wang, Yan & Wang, Yuanchao (2018) Trick or Treat: Microbial Pathogens Evolved Apoplastic Effectors Modulating Plant Susceptibility to Infection. Molecular Plant-Microbe Interactions®, 31, 6–12. 10.1094/MPMI-07-17-0177-FI.

Watson, B.S., Bedair, M.F., Urbanczyk-Wochniak, E., Huhman, D.V., Yang, D.S., Allen, S.N., et al. (2015) Integrated metabolomics and transcriptomics reveal enhanced specialized metabolism in Medicago truncatula root border cells. Plant Physiology, 167, 1699–1716. 10.1104/pp.114.253054.

White, F., Potnis, N., Jones, J.B. & Koebnik, R. (2009) The type III effectors of Xanthomonas. Molecular Plant Pathology, 10, 749–766. 10.1111/j.1364-3703.2009.00590.x.

Wu, T., Zhang, H., Yuan, B., Liu, H., Kong, L., Chu, Z., et al. (2022) Tal2b targets and activates the expression of OsF3H03g to hijack OsUGT74H4 and synergistically interfere with rice immunity. New Phytologist, 233, 1864–1880. 10.1111/nph.17877.

Yan, D., Tajima, H., Cline, L.C., Fong, R.Y., Ottaviani, J.I., Shapiro, H.-Y., et al. (2022) Genetic modification of flavone biosynthesis in rice enhances biofilm formation of soil diazotrophic bacteria and biological nitrogen fixation. Plant Biotechnology Journal, 20, 2135–2148. 10.1111/pbi.13894.

Yang, B., Sugio, A. & White, F.F. (2006) Os8N3 is a host disease-susceptibility gene for bacterial blight of rice. Proc Natl Acad Sci U S A, 103, 10503–10508. 10.1073/pnas.0604088103.

Yang, Z., Li, N., Kitano, T., Li, P., Spindel, J.E., Wang, L., et al. (2021) Genetic mapping identifies a rice naringenin O-glucosyltransferase that influences insect resistance. The Plant Journal, 106, 1401–1413. 10.1111/tpj.15244.

Yu, P., He, X., Baer, M., Beirinckx, S., Tian, T., Moya, Y.A.T., et al. (2021) Plant flavones enrich rhizosphere Oxalobacteraceae to improve maize performance under nitrogen deprivation. Nature Plants, 7, 481–499. 10.1038/s41477-021-00897-y.

Zeilmaker, T., Ludwig, N.R., Elberse, J., Seidl, M.F., Berke, L., Van Doorn, A., et al. (2015) DOWNY MILDEW RESISTANT 6 and DMR 6-LIKE OXYGENASE 1 are partially redundant but distinct suppressors of immunity in Arabidopsis. The Plant Journal, 81, 210–222. 10.1111/tpj.12719.

Zhang, H., Zhang, F., Yu, Y., Feng, L., Jia, J., Liu, B., et al. (2020) A Comprehensive Online Database for Exploring ∼20,000 Public Arabidopsis RNA-Seq Libraries. Molecular Plant, 13, 1231–1233. 10.1016/j.molp.2020.08.001.

Zhang, Y., Yu, Q., Gao, S., Yu, N., Zhao, L., Wang, J., et al. (2022) Disruption of the primary salicylic acid hydroxylases in rice enhances broad-spectrum resistance against pathogens. *Plant*, Cell & Environment, 45, 2211–2225. 10.1111/pce.14328.

Zhang, Y., Zhao, L., Zhao, J., Li, Y., Wang, J., Guo, R., et al. (2017) *S5H/DMR6* Encodes a Salicylic Acid 5-Hydroxylase That Fine-Tunes Salicylic Acid Homeostasis. Plant Physiology, 175, 1082–1093. 10.1104/pp.17.00695.

Zhu, Y.-X., Ge, C., Ma, S., Liu, X.-Y., Liu, M., Sun, Y., et al. (2020) Maize ZmFNSI Homologs Interact with an NLR Protein to Modulate Hypersensitive Response. International Journal of Molecular Sciences, 21, 2529. 10.3390/ijms21072529.

