## Supplementary material for "*Xanthomonas oryzae* pv. *oryzae* type-III effector TAL9b targets a broadly conserved disease susceptibility locus to promote pathogenesis in rice": Figure S

[†] Equally contributed authors

[‡] Corresponding authors

### Supplementary Figures

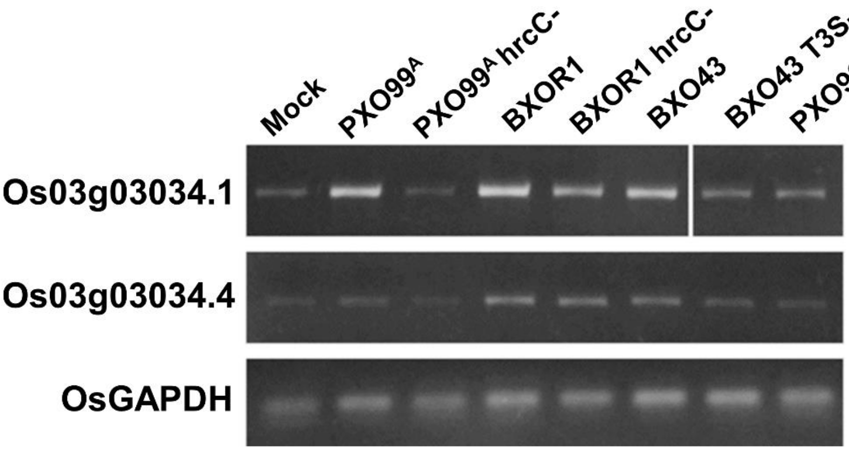


**Figure S1: The splice variant *LOC_Os03g03034.1* is primarily induced by Xoo and Xoc.** Bacterial suspensions of indicated strains (OD$_{600}$ = 1) were infiltrated into the leaves of 14-day-old rice leaves, sampled at 24 hours post infiltration and the gene expression was analysed through semi-quantitative PCR using the transcript variant specific primer pairs. *OsGAPDH* was used as an internal control for equal input RNA.

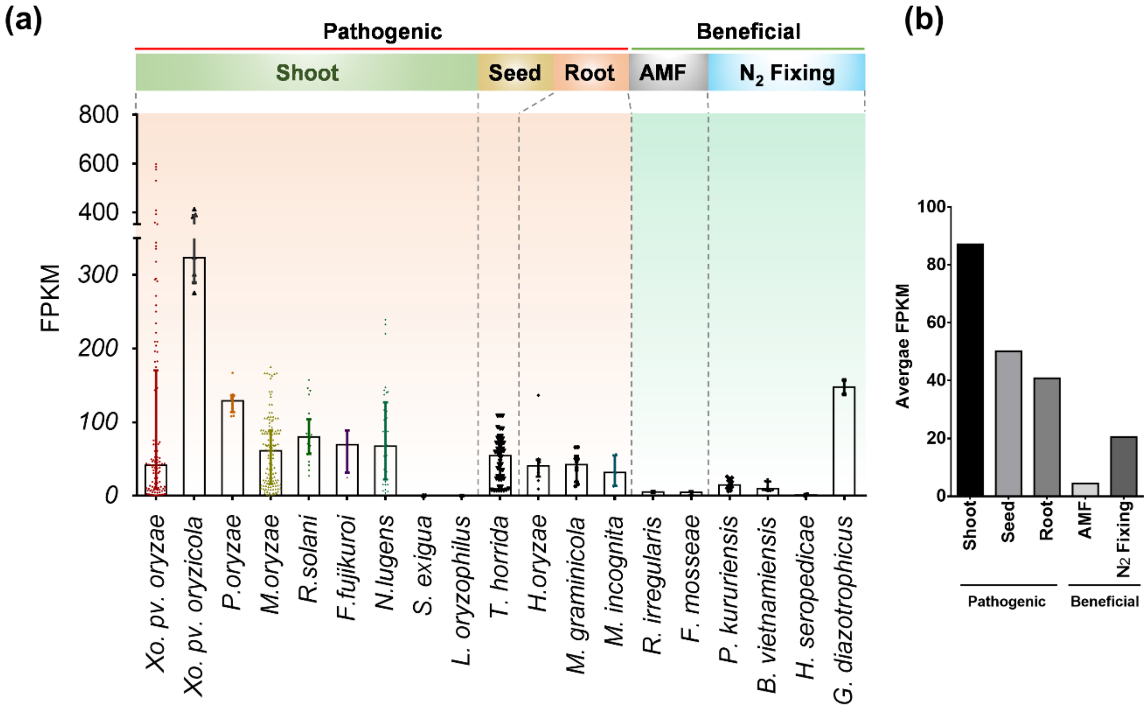


**Figure S2: Expression of *LOC_Os03g03034.1* is induced majorly by foliar/shoot pathogens.** Normalised expression values of *LOC_Os03g03034.1* during rice-biotic factor interaction were obtained from the Rice RNA-seq database (http://ipf.sustech.edu.cn/pub/ricerna/) and categorised based on tissue and nature of the interaction. (a) Bar graph showing the expression values (FPKM) of the gene during the interaction between rice and individual pathogenic or beneficial organisms. (b) Average FPKM values based on tissue and interaction type were calculated using values plotted in (a).

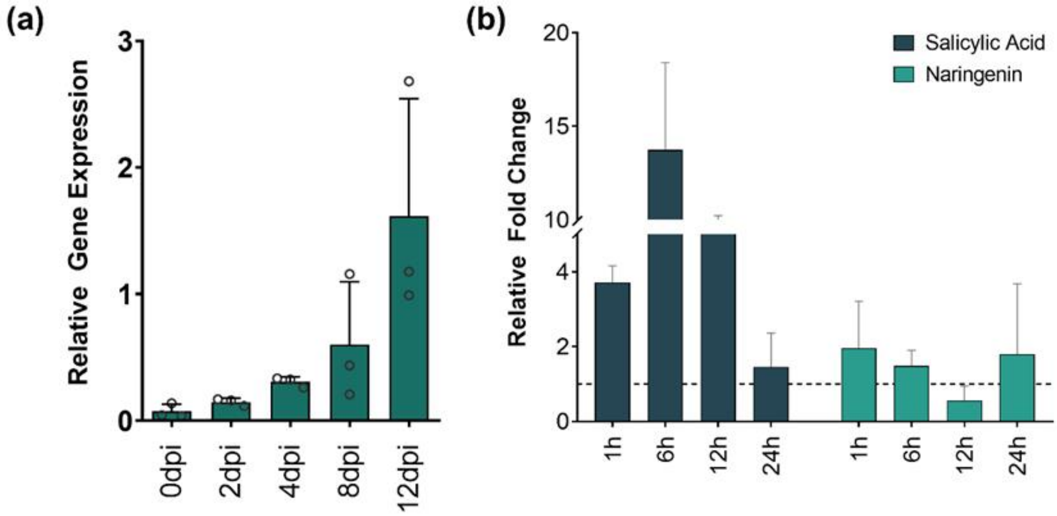


**Figure S3: *OsS5H/FNS-03g* is induced dynamically by Xoo, SA and Naringenin.** Expression of *OsS5H/FNS-03g* in rice leaves was assessed using qRT-PCR at the indicated time points post Xoo PXO99$^{A}$ clip inoculation (a) or post exogenous application of SA or Naringenin (b). The gene expression levels were normalised with the internal control *OsGAPDH*. Relative gene expression in (a) was calculated using the $2^{-\Delta Ct}$ method. Fold change values in (b) were calculated using the $2^{-\Delta\Delta Ct}$ method relative to the mock-treated samples at respective timepoints. Error bars in (a) and (b) indicate the standard deviation of the gene expression levels and fold change values, respectively, in three independent experimental replicates.

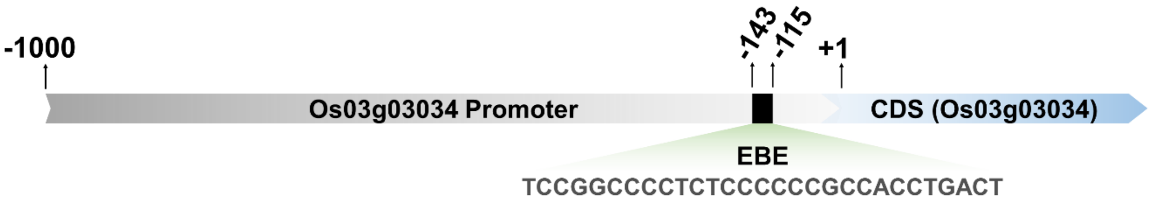


**Figure S4: *LOC_Os03g03034* promoter harbours an effector binding element (EBE).** Graphical representation of the location of the EBE in the upstream sequence of *LOC_Os03g03034*. +1 indicates the start codon and the numbers prefixed with "-" indicate the position in the promoter of the gene relative to the start codon. CDS - Coding sequence.

Xoo

| TALE by strain | RVD | | | | | | | | | | | | | | | | | | | | | | | | | |
|---|---|---|---|---|---|---|---|---|---|---|---|---|---|---|---|---|---|---|---|---|---|---|---|---|---|---|---|
| TalAQ15 Xoo PXO142 (Pseudo) | HD | HD | NN | NN | NI | NG | HD | S* | HG | HD | NG | N* | NG | HD | – | – | – | – | – | – | – | – | – | – | – | – | – |
| TalAQ55 Xoo IXO842 | HD | HD | NN | NN | NI | NG | HD | S* | HG | HD | NG | N* | NG | HD | HD | N* | NI | NI | NI | HD | HI | ND | HD | NG | NN | HG | N* |
| TalAQ29 Xoo SK2-3 | HD | HD | NN | NN | NI | NG | HD | S* | HG | HD | NG | N* | NG | HD | HD | N* | NI | NI | NN | HD | HI | ND | HD | NG | NN | HG | N* |
| TalAQ44 Xoo IXO221 | HD | HD | NN | NN | NI | NG | HD | S* | HG | HD | NG | N* | NG | HD | HD | N* | NI | NI | NN | HD | HI | ND | HD | NG | NN | HG | N* |
| TalAQ32 Xoo IXO704 | HD | HD | NN | NN | NI | NG | HD | S* | HG | HD | NG | N* | NG | HD | HD | N* | NI | NI | NN | HD | HI | ND | HD | NG | NN | HG | N* |
| TalAQ46 Xoo IXO493 | HD | HD | NN | NN | NI | NG | HD | S* | HG | HD | NG | N* | NG | HD | HD | N* | NI | NI | NN | HD | HI | ND | HD | NG | NN | HG | N* |
| TalAQ49 Xoo IXO630 | HD | HD | NN | NN | NI | NG | HD | S* | HG | HD | NG | N* | NG | HD | HD | N* | NI | NI | NN | HD | HI | ND | HD | NG | NN | HG | N* |
| TalAQ60 Xoo PXO35 | HD | HD | NN | NN | NI | NG | HD | S* | HG | HD | NG | N* | NG | HD | HD | N* | NI | NI | NN | HD | HI | ND | HD | NG | NN | HG | N* |
| TalAQ24 Xoo PXO421 | HD | HD | NN | NN | NI | NG | HD | S* | HG | HD | NG | N* | NG | HD | HD | N* | NI | NI | NN | HD | HI | ND | HD | NG | NN | HG | N* |
| TalAQ13 Xoo XF89b | HD | HD | NN | NN | NI | NG | HD | S* | HG | HD | NG | N* | NG | HD | HD | N* | NI | NI | NN | HD | HI | ND | HD | NG | NN | HG | N* |
| TalAQ41 Xoo IXO98 | HD | HD | NN | NN | NI | NG | HD | S* | HG | HD | NG | N* | NG | HD | HD | N* | NI | NI | NN | HD | HI | ND | HD | NG | NN | HG | N* |
| TalAQ11 Xoo PXO602 | HD | HD | NN | NN | NI | NG | HD | S* | HG | HD | NG | N* | NG | HD | HD | N* | NI | NI | NN | HD | HI | ND | HD | NG | NN | HG | N* |
| TalAQ54 Xoo IXO792 | HD | HD | NN | NN | NI | NG | HD | S* | HG | HD | NG | N* | NG | HD | HD | N* | NI | NI | NN | HD | HI | ND | HD | NG | NN | HG | N* |
| TalAQ21 Xoo JW11089 | HD | HD | NN | NN | NI | NG | HD | S* | HG | HD | NG | N* | NG | HD | HD | N* | NI | NI | NN | HD | HI | ND | HD | NG | NN | HG | N* |
| TalAQ16 Xoo AUST2013 | HD | HD | NN | NN | NI | NG | HD | S* | HG | HD | NG | N* | NG | HD | HD | N* | NI | NI | NN | HD | HI | ND | HD | NG | NN | HG | N* |
| TalAQ9 Xoo PXO524 | HD | HD | NN | NN | NI | NG | HD | S* | HG | HD | NG | N* | NG | HD | HD | N* | NI | NI | NN | HD | HI | ND | HD | NG | NN | HG | N* |
| TalAQ42 Xoo IXO134 | HD | HD | NN | NN | NI | NG | HD | S* | HG | HD | NG | N* | NG | HD | HD | N* | NI | NI | NN | HD | HI | ND | HD | NG | NN | HG | N* |
| TalAQ50 Xoo IXO644 | HD | HD | NN | NN | NI | NG | HD | S* | HG | HD | NG | N* | NG | HD | HD | N* | NI | NI | NN | HD | HI | ND | HD | NG | NN | HG | N* |
| TalAQ52 Xoo IXO675 (Pseudo) | HD | HD | NN | NN | NI | NG | HD | S* | HG | HD | NG | N* | NG | HD | HD | N* | NI | NI | NN | HD | HI | ND | HD | NG | NN | HG | N* |
| TalAQ65 Xoo CR1-5 | HD | HD | NN | NN | NI | NG | HD | S* | HG | HD | NG | N* | NG | HD | HD | N* | NI | NI | NN | HD | HI | ND | HD | NG | NN | HG | N* |
| TalAQ25 Xoo PXO513 | HD | HD | NN | NN | NI | NG | HD | S* | HG | HD | NG | N* | NG | HD | HD | N* | NI | NI | NN | HD | HI | ND | HD | NG | NN | HG | N* |
| TalAQ63 Xoo GXO2001 | HD | HD | NN | NN | NI | NG | HD | S* | HG | HD | NG | N* | NG | HD | HD | N* | NI | NI | NN | HD | HI | ND | HD | NG | NN | HG | N* |
| TalAQ34 Xoo K1 | HD | HD | NN | NN | NI | NG | HD | S* | HG | HD | NG | N* | NG | HD | HD | N* | NI | NI | NN | HD | HI | ND | HD | NG | NN | HG | N* |
| TalAQ5 Xoo PXO71 | HD | HD | NN | NN | NI | NG | HD | S* | HG | HD | NG | N* | NG | HD | HD | N* | NI | NI | NN | HD | HI | ND | HD | NG | NN | HG | N* |
| TalAQ22 Xoo KXO85 | HD | HD | NN | NN | NI | NG | HD | S* | HG | HD | NG | N* | NG | HD | HD | N* | NI | NI | NN | HD | HI | ND | HD | NG | NN | HG | N* |
| TalAQ64 Xoo GXO2016 | HD | HD | NN | NN | NI | NG | HD | S* | HG | HD | NG | N* | NG | HD | HD | N* | NI | NI | NN | HD | HI | ND | HD | NG | NN | HG | N* |
| TalAQ43 Xoo IXO220 | HD | HD | NN | NN | NI | NG | HD | S* | HG | HD | NG | N* | NG | HD | HD | N* | NI | NI | NN | HD | HI | ND | HD | NG | NN | HG | N* |
| TalAQ35 Xoo K2 | HD | HD | NN | NN | NI | NG | HD | S* | HG | HD | NG | N* | NG | HD | HD | N* | NI | NI | NN | HD | HI | ND | HD | NG | NN | HG | N* |
| TalAQ26 Xoo PXO61 | HD | HD | NN | NN | NI | NG | HD | S* | HG | HD | NG | N* | NG | HD | HD | N* | NI | NI | NN | HD | HI | ND | HD | NG | NN | HG | N* |
| TalAQ39 Xoo IXO74 | HD | HD | NN | NN | NI | NG | HD | S* | HG | HD | NG | N* | NG | HD | HD | N* | NI | NI | NN | HD | HI | ND | HD | NG | NN | HG | N* |
| TalAQ45 Xoo IXO365 | HD | HD | NN | NN | NI | NG | HD | S* | HG | HD | NG | N* | NG | HD | HD | N* | NI | NI | NN | HD | HI | ND | HD | NG | NN | HG | N* |
| TalAQ38 Xoo IXO35 | HD | HD | NN | NN | NI | NG | HD | S* | HG | HD | NG | N* | NG | HD | HD | N* | NI | NI | NN | HD | HI | ND | HD | NG | NN | HG | N* |
| TalAQ66 Xoo FY517 | HD | HD | NN | NN | NI | NG | HD | S* | HG | HD | NG | N* | NG | HD | HD | N* | NI | NI | NN | HD | HI | ND | HD | NG | NN | HG | N* |
| TalAQ23 Xoo PXO404 | HD | HD | NN | NN | NI | NG | HD | S* | HG | HD | NG | N* | NG | HD | HD | N* | NI | NI | NN | HD | HI | ND | HD | NG | NN | HG | N* |
| TalAQ57 Xoo T7133 | HD | HD | NN | NN | NI | NG | HD | S* | HG | HD | NG | N* | NG | HD | HD | N* | NI | NI | NN | HD | HI | ND | HD | NG | NN | HG | N* |
| TalAQ62 Xoo GXO2006 | HD | HD | NN | NN | NI | NG | HD | S* | HG | HD | NG | N* | NG | HD | HD | N* | NI | NI | NN | HD | HI | ND | HD | NG | NN | HG | N* |
| TalAQ17 Xoo IX-280 | HD | HD | NN | NN | NI | NG | HD | S* | HG | HD | NG | N* | NG | HD | HD | N* | NI | NI | NN | HD | HI | ND | HD | NG | NN | HG | N* |
| TalAQ12 Xoo PXO602 (Pseudo) | – | – | – | – | – | – | – | – | HG | HD | NG | N* | NG | HD | HD | N* | NI | NI | NN | HD | HI | ND | HD | NG | NN | HG | N* |
| TalAQ33 Xoo IXO1088 | HD | HD | NN | NN | NG | NG | HD | NS | HG | HD | NG | N* | HD | HD | HD | N* | NN | NI | NN | HD | HI | ND | HD | HG | NN | HG | N* |
| TalAQ27 Xoo PXO79 | HD | HD | NN | NN | NG | NG | HD | NS | HG | HD | NG | N* | HD | HD | HD | N* | NN | NI | NN | HD | HI | ND | HD | HG | NN | HG | N* |
| TalAQ14 Xoo ICMP3125 | HD | HD | NN | NN | NG | NG | HD | NS | HG | HD | NG | N* | HD | HD | HD | N* | NN | NI | NN | HD | HI | ND | HD | HG | NN | HG | N* |
| TalAQ56 Xoo IXO1104 | HD | HD | NN | NN | NG | NG | HD | NS | HG | HD | NG | N* | HD | HD | HD | N* | NN | NI | NN | HD | HI | ND | HD | HG | NN | HG | N* |
| TalAQ1 Xoo PXO99A (tal9b) | HD | HD | NN | NN | NG | NG | HD | NS | HG | HD | NG | N* | HD | HD | HD | N* | NN | NI | NN | HD | HI | ND | HD | HG | NN | HG | N* |
| TalAQ59 Xoo FXO38 | HD | HD | NN | NN | NG | NG | HD | NS | HG | HD | NG | HD | HD | HD | HD | N* | NN | NI | NN | HD | HI | ND | HD | HG | NN | HG | N* |
| TalAQ51 Xoo IXO651 | HD | HD | NN | NN | NN | NG | HD | S* | HG | HD | NG | N* | HD | HD | HD | N* | NN | NI | NN | HD | HI | ND | HD | HG | NI | HG | N* |
| TalAQ6 Xoo PXO145 | HD | HD | NN | NN | NS | NG | HD | S* | HG | HD | NG | N* | HD | HD | HD | N* | NN | NI | NN | HD | HM | ND | HD | HG | AN | HG | N* |
| TalAQ48 Xoo IXO608 | HD | HD | NN | NN | NS | NG | HD | S* | HG | HD | NG | N* | HD | HD | HD | N* | NN | NI | NN | HD | HI | ND | HD | HG | NN | HG | N* |
| TalAQ40 Xoo IXO97 | HD | HD | NN | NN | NS | NG | HD | S* | HG | HD | NG | N* | HD | HD | HD | N* | NN | NI | NN | HD | HI | ND | HD | HG | NN | HG | N* |
| TalAQ18 Xoo JL25 | HD | HD | NN | NN | NS | NG | HD | S* | HG | HD | NG | N* | HD | HD | HD | N* | NN | NI | NN | HD | HI | ND | HD | HG | NN | HG | N* |
| TalAQ61 Xoo Huang604 | HD | HD | NN | NN | NS | NG | HD | S* | HG | HD | NG | N* | HD | HD | HD | N* | NN | NI | NN | HD | HI | ND | HD | HG | NN | HG | N* |
| TalAQ10 Xoo PXO563 | HD | HD | NN | NN | NS | NG | HD | S* | HG | HD | NG | N* | HD | HD | HD | N* | NN | NI | NN | HD | HI | ND | HD | HG | NN | HG | N* |
| TalAQ36 Xoo K3 | HD | HD | NN | NN | NS | NG | HD | S* | HG | HD | NG | N* | HD | HD | HD | N* | NN | NI | NN | HD | HI | ND | HD | HG | NN | HG | N* |
| TalAQ7 Xoo PXO211 | HD | HD | NN | NN | NS | NG | HD | S* | HG | HD | NG | N* | HD | HD | HD | N* | NN | NI | NN | HD | HI | ND | HD | HG | NN | HG | N* |
| TalAQ47 Xoo IXO597 | HD | HD | NN | NN | NS | NG | HD | S* | HG | HD | NG | N* | HD | HD | HD | N* | NN | NI | NN | HD | HI | ND | HD | HG | NN | HG | N* |
| TalAQ2 Xoo MAFF311018 (XOO1136) | HD | HD | NN | NN | NS | NG | HD | S* | HG | HD | NG | N* | HD | HD | HD | N* | NN | NI | NN | HD | HI | ND | HD | HG | NN | HG | N* |
| TalAQ37 Xoo K3a | HD | HD | NN | NN | NS | NG | HD | S* | HG | HD | NG | N* | HD | HD | HD | N* | NN | NI | NN | HD | HI | ND | HD | HG | NN | HG | N* |
| TalAQ3 Xoo PXO83 | HD | HD | NN | NN | NS | NG | HD | S* | HG | HD | NG | N* | HD | HD | HD | N* | NN | NI | NN | HD | HI | ND | HD | HG | NN | HG | N* |
| TalAQ28 Xoo ScYc-b | HD | HD | NN | NN | NS | NG | HD | S* | HG | HD | NG | N* | HD | HD | HD | N* | NN | NI | NN | HD | HI | ND | HD | HG | NN | HG | N* |
| TalAQ30 Xoo XM9 | HD | HD | NN | NN | NS | NG | HD | S* | HG | HD | NG | N* | HD | HD | HD | N* | NN | NI | NN | HD | HI | ND | HD | HG | NN | HG | N* |
| TalAQ31 Xoo YC11 | HD | HD | NN | NN | NS | NG | HD | S* | HG | HD | NG | N* | HD | HD | HD | N* | NN | NI | NN | HD | HI | ND | HD | HG | NN | HG | N* |
| TalAQ4 Xoo PXO86 (tal8b) | HD | HD | NN | NN | NS | NG | HD | S* | HG | HD | NG | N* | HD | HD | HD | N* | NN | NI | NN | HD | HI | ND | HD | HG | NN | HG | N* |
| TalAQ53 Xoo IXO685 | HD | HD | NN | NN | NS | NG | HD | S* | HG | HD | NG | N* | HD | HD | HD | N* | NN | NI | NN | HD | HI | ND | HD | HG | NN | HG | N* |
| TalAQ8 Xoo PXO236 | HD | HD | NN | NN | NS | NG | HD | S* | HG | HD | NG | N* | HD | HD | HD | N* | NN | NI | NN | HD | HI | ND | HD | HG | NN | HG | N* |
| TalAQ20 Xoo JP01 | HD | HD | NN | NN | NS | NG | HD | S* | HG | HD | NG | N* | HD | HD | HD | N* | NN | NI | NN | HD | HI | ND | HD | HG | NN | HG | N* |
| TalAQ58 Xo AH28 | – | HD | HD | NN | NS | NG | HD | S* | HG | HD | NG | N* | HD | HD | HD | N* | NN | NI | NN | HD | HI | ND | HD | HG | NN | HG | N* |
| TalAQ19 Xoo JL28 | QY | G* | QY | NN | NS | NG | HD | S* | HG | HD | NG | N* | HD | HD | HD | N* | NN | NI | NN | HD | HI | ND | HD | HG | NN | HG | N* |
| LOC_Os03g03034 EBE TC | C | G | G | C | C | C | C | T | C | T | C | C | C | C | C | C | G | C | C | A | C | C | T | G | A | C | T |

Xoc

| TALE by strain | RVD | | | | | | | | | | | | | | | | | | | | | | | | | |
|---|---|---|---|---|---|---|---|---|---|---|---|---|---|---|---|---|---|---|---|---|---|---|---|---|---|---|---|
| TalBR3 Xoc BLS279 | HD | HD | NN | NN | NG | N* | HD | NI | NG | HD | NG | NS | ND | HA | ND | N* | ND | NN | HD | NN | NN | HD | HD | N* | NN | NG | HD |
| TalBR1 Xoc BLS256 | HD | HD | NN | NN | NG | N* | HD | NI | NG | HD | NG | NS | HD | HA | ND | N* | ND | NN | HD | NN | NN | HD | HD | N* | NN | NG | HD |
| TalBR5 Xoc CFBP2286 | HD | HD | NN | NN | NG | N* | HD | NI | NG | HD | NG | NS | HD | HA | ND | N* | ND | NN | HD | NN | NN | HD | HD | N* | NN | NG | HD |
| TalBR2 Xoc B8-12 | HD | HD | NN | NN | NG | N* | HD | NI | NG | HD | NG | NS | HD | HA | HD | N* | HD | NN | HD | NN | NN | HD | HD | NG | NN | NG | HD |
| TalBR6 Xoc L8 | HD | HD | NN | NN | NG | N* | HD | NI | NG | HD | NG | NS | HD | HA | HD | N* | HD | NN | HD | NN | NN | HD | HD | NG | NN | NG | HD |
| TalBR9 Xoo HGA4 | HD | HD | NN | NN | NG | N* | HD | NI | NG | HD | NG | NS | HD | HA | HD | N* | HD | NN | HD | NN | NN | HD | HD | NG | NN | NG | HD |
| TalBR8 Xoc 0-9 | HD | HD | NN | NN | NG | N* | HD | NI | NG | HD | NG | NS | ND | HA | HD | N* | HD | NG | HD | NN | NN | HD | HD | NG | NN | NG | HD |
| TalBR4 Xoc BXOR1 | HD | HD | NN | NN | NG | N* | HD | NI | NG | HD | NG | NS | ND | N* | HD | N* | HD | NG | HD | NN | NN | HD | HD | NG | NN | NG | HD |
| TalBR10 Xoc YN01 | HD | HD | NN | NN | NG | N* | HD | NI | NG | HD | NG | NS | ND | N* | HD | N* | HD | NG | HD | NN | NN | HD | HD | NG | NN | NG | HD |
| TalBR7 Xoc GX01 | HD | HD | NN | NN | NG | N* | HD | NI | NG | HD | NG | NS | ND | N* | HD | N* | HD | NG | HD | NN | NN | HD | HD | NG | NN | NG | HD |
| TalBR11 Xoc ML5 | HD | HD | NN | NN | NG | N* | HD | NI | NG | HD | NG | NS | ND | N* | HD | N* | HD | NG | HD | NN | NN | HD | HD | NG | NN | NG | HD |
| LOC_Os03g03034 EBE TC | C | G | G | C | C | C | C | T | C | T | C | C | C | C | C | C | G | C | C | A | C | C | T | G | A | C | T |

**Figure S5: Majority of the Xoo and Xoc strains encode TALEs that induce *LOC_Os03g03034*.** Shown here are Repeat Variable Diresidues (RVDs) of the TALEs, from various Xoo and Xoc strains, belonging to the classes that were shown to induce the expression of *LOC_Os03g03034*. Each RVD type is differently coloured.

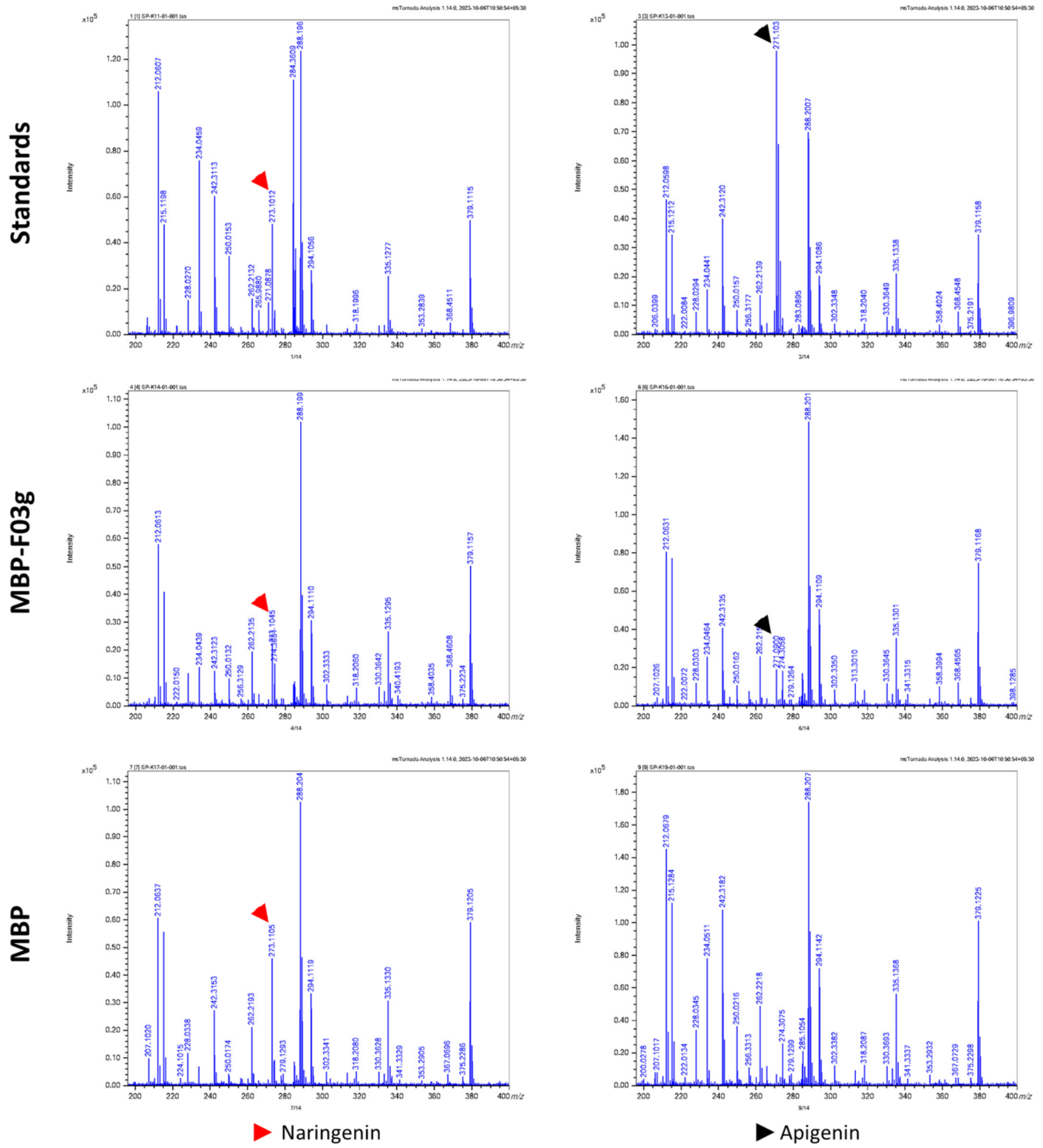


**Figure S6: OsS5H/FNS-03g catalyses the formation of apigenin from naringenin.** MALDI-TOF spectrum of fractions collected from the standards (pure naringenin and apigenin), biochemical assay mixture containing MBP-F03g, and MBP alone. The fractions were collected at the retention time when naringenin and apigenin were detected. The red arrowhead indicates the peak corresponding to Naringenin (m/z=273) and the black arrowhead indicates the peak corresponding to Apigenin (m/z=271). MBP - Maltose Binding Protein; MBP-F03g - MBP-OsS5H/FNS-03g fusion protein.

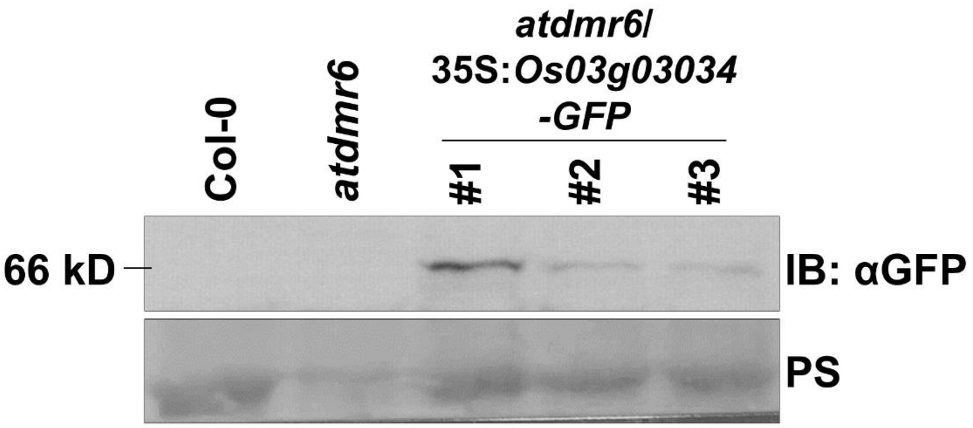


**Figure S7: Confirmation of LOC_Os03g03034.1-GFP protein expression in *Arabidopsis atdmr6* complementation lines.** Protein extracts from the leaves from *Arabidopsis atdmr6* transgenic plants (three independent lines) expressing Os03g03034-GFP (MW = ~66.7kD) were subjected to immunoblotting using an anti-GFP antibody (top panel). Total protein was visualised using Ponceau S staining (PS; bottom panel).

### Supplementary Tables

**Table S1**: List of bacterial strains, plant materials, and plasmids used in this study.

| S. No. | Organism[a] | Strain | Construct/Genotype | Source | Selection[b] |
|---|---|---|---|---|---|
| **List of Bacterial materials** | | | | | |
| 1 | *E.coli* | DH5α | pK18mob:PXO99A hrcC | This study | Kan |
| 2 | *E.coli* | DH5α | pK18mob:BXORI hrcC | This study | Kan |
| 3 | *E.coli* | DH5α | pK18mob:PXO99A tal9b | This study | Kan |
| 4 | *E.coli* | S17-1 | pK18mob:PXO99A hrcC | This study | Kan |
| 5 | *E.coli* | S17-1 | pK18mob:BXORI hrcC | This study | Kan |
| 6 | *E.coli* | S17-1 | pK18mob:PXO99A tal9b | This study | Kan |
| 7 | *E.coli* | DH5α | pENTR:LOC_Os03g03034.1 | This study | Kan |
| 8 | *E.coli* | DH5α | pENTR:LOC_Os03g03034.1 | This study | Kan |
| 9 | *E.coli* | DH5α | pH7FWG2:LOC_Os03g03034.1 | This study | Spec |
| 10 | *E.coli* | DH5α | pETM-40:LOC_Os03g03034.1 | This study | Kan |
| 11 | *E.coli* | DH5α | pETM-40 | This study | Kan |
| 12 | *E.coli* | Rosetta DE3 | pETM-40:LOC_Os03g03034.1 | This study | Kan; Chlr |
| 13 | *E.coli* | Rosetta DE3 | pETM-40 | This study | Kan; Chlr |
| 14 | *A.tumefaciens* | AGL1 | pH7FWG2:LOC_Os03g03034.1 | This study | Rif; Carb; Spec |
| 15 | Xoo | PXO99$^{A}$ | Wildtype | Lab stock | Rif |
| 16 | Xoo | PXO86 | Wildtype | Lab stock | Rif |
| 17 | Xoo | BXO43 | Wildtype | Lab stock | Rif |
| 18 | Xoo | PXO99$^{A}$ hrcC- | hrcC mutant (T3SS deficient) | This study | Rif; Kan |
| 19 | Xoo | PXO99$^{A}$ tal9b- | tal9b mutant | This study | Rif; Kan |
| 20 | Xoo | BXO43 t3s- | T3S deficient | Lab stock | Rif; Carb |
| 21 | Xoc | BLS256 | Wildtype | Lab stock | Rif |
| 22 | Xoc | BXORI | Wildtype | Lab stock | Rif |
| 23 | Xoc | BXORI hrcC- | hrcC mutant (T3SS deficient) | This study | Rif; Kan |
| 24 | Pst | DC3000 | Wildtype | Lab stock | Rif |
| **List of Plant materials** | | | | | |
| 24 | *O. sativa* | TN1 | Wildtype; bacterial blight susceptible | Lab stock | |
| 25 | *A. thaliana* | Col-0 | Wildtype | Lab stock | |
| 26 | *A. thaliana* | *atdmr6* (SK19087) | Null mutant of *AtDMR6* | Procured | Phosphinothricin |
| 27 | *A. thaliana* | *atdmr6*/pH7FWG2: *LOC_Os03g03034.1* | A null mutant of *AtDMR6* ectopically expressing *LOC_Os03g03034-eGFP* | This study | Phosphinothricin; Hygromycin |

[a] *E.coli - Escherichia coli*; *A.tumefaciens - Agrobacterium tumefaciens*; Xoo - *Xanthomonas oryzae* pv. *oryzae;* Xoc - *Xanthomonas oryzae* pv. *oryzicola;* Pst - *Pseudomonas syringae* pv. *tomato*; *O. sativa - Oryza sativa*; *A. thaliana - Arabidopsis thaliana*

[b] Kan - Kanamycin; Spec - Spectinomycin; Rif - Rifampicin; Carb - Carbenicillin; Chlr - Chloramphenicol

**Table S2**: List of publicly available microarray data analysed in this study.

| S.No. | NCBI GEO ID | Pathogen[a] | Time point[b] |
|---|---|---|---|
| 1 | GSE36272 | Xoo | 24hpi |
| 2 | GSE33411 | Xoo | 24hpi |
| 3 | GSE16793 | Xoo and Xoc | 24hpi |
| 4 | GSE7256 | *M. oryzae* | 72hpi and 96hpi |
| 5 | GSE95394 | *M. oryzae* | 48hpi and 72hpi |
| 6 | GSE28308 | *M. oryzae* | 72hpi |
| 7 | GSE880246 | Bph | 3hpi and 6hpi |
| 8 | GSE30707 | *Puccinia triticina* f. sp. *Tritici* | 24hpi |

[a] Xoo - *Xanthomonas oryzae* pv. *oryzae;* Xoc - *Xanthomonas oryzae* pv. *oryzicola; M. oryzae - Magnaporthe oryzae;* Bph - *Brown plant hopper.*

[b] hpi - hours post infection

**Table S3**: List of all the primers used in this study.

| S.No. | Primer | Sequence (5'-3') | Purpose |
|---|---|---|---|
| 1 | LOC_Os03g03034.1_F | caccATGGCGGACCAGCTCATCTC | Primers for gene cloning and semi-quantitative PCR |
| 2 | LOC_Os03g03034.4_F | caccATGCAATCGATCGATGGAGT | |
| 3 | LOC_Os03g03034_R | AGATGTGTCTGTAGGTGTTGTT | |
| 4 | LOC_Os03g03034.1_RT_F | TTCTCAAGGAAGGCAGGTGGATCG | qRT-PCR primers |
| 5 | LOC_Os03g03034.1_RT_R | TTCCGTTGCTTAGCGCCTGTAG | |
| 6 | OsGAPDH_RT_F | AAGCCAGCATCCTATGATCAGATT | |
| 7 | OsGAPDH_RT_R | CGTAACCCAGAATACCCTTGAGTTT | |
| 8 | 99A-R1_hrcC_EcoRI-F | ggaattcTGCAGATTCCAGGCATGGCC | Primers for amplifying homologous fragment for knocking out hrcC |
| 9 | 99A-R1_hrcC_BamH1-R | cgggatccGATGATGGTGGCATCGATCTGCA | |
| 10 | ICF_99A_hrcC | CGGCCATTGCCAAGCAGGTC | Primers to confirm vector integration |
| 11 | ICF_R1_hrcC | CGGCGATTGCCAGGCAGGTC | |
| 12 | ICR_99A-R1_hrcC | GCTCAATACGCCGTTGAAACCCA | |
| 13 | Tal9b_INS_EcoRI-F | ggaattcCGACACACGAAGACATCGTTGGC | Primers for amplifying homologous fragment for knocking out Tal9b |
| 14 | Tal9b_INS_BamH1-R | cgggatccGATGGCCACCGCCTGCG | |
| 15 | Tal9b-ICF | ACTGGTGGGCCATGGGTTTAC | Primers to confirm vector integration |
| 16 | Tal9b-ICR | ATTGCTGGCGATGGCCACCAC | |
| 17 | OsF3-RFC_F | ATTTTCAGGGCGCCATGGACATGGCGGACCAGCTCATCTC | Primers to clone *LOC_Os03g03034* in pETM-40 using restriction-free cloning |
| 18 | OsF3-RFC_R | TCAGTGGTGGTGGTGGTGGTGAGATGTGTCTGTAGGTGTTG | |
| 19 | pSKTAIL-L1_F | TTCTCATCTAAGCCCCCATTTGG | Primers to genotype *atdmr6* mutants |
| 20 | DMR6-genomic-F | CCTCCATGTCCTGAACCTGAGC | |
| 21 | DMR6-genomic-R | CGAGGCAATGTTCTTGGTCCAG | |
